# Repetitive Levodopa Treatment Drives Cell Type-Specific Striatal Adaptations Associated with Progressive Dyskinesia in Parkinsonian Mice

**DOI:** 10.1101/2025.05.16.654598

**Authors:** Rodrigo Manuel Paz, Michael B. Ryan, Pamela F. Marcott, Allison E. Girasole, Joe Faryean, Vincent Duong, Sadhana Sridhar, Alexandra B. Nelson

## Abstract

The use of levodopa to manage Parkinson’s disease (PD) symptoms leads to levodopa-induced dyskinesia (LID) and other motor fluctuations, which worsen with disease progression and repeated treatment. Aberrant activity of striatal D1- and D2-expressing medium spiny neurons (D1-/D2-MSNs) underlies LID, but the mechanisms driving its progression remain unclear. Using the 6-OHDA mouse model of PD/LID, we combined *in vivo* and *ex vivo* recordings to isolate the effect of repeated treatment in LID worsening and other changes in the motor response to levodopa. We found that LID worsening is linked to potentiation of levodopa-evoked responses in both D1-/D2-MSNs, independent of changes in dopamine release or MSN intrinsic excitability. Instead, strengthening of glutamatergic synapses onto D1-MSNs emerged as a key signature. Moreover, we found changes in D2-MSN activity are associated with LID duration, which paralleled a reduction in D2R sensitivity. These findings reveal striatal adaptations contributing to worsening of levodopa-related complications.

## INTRODUCTION

In Parkinson’s disease (PD), the degeneration of dopamine neurons in the substantia nigra pars compacta leads to cardinal motor symptoms, including bradykinesia, rigidity and tremor^1,2^. Dopamine replacement therapy with levodopa remains the most effective symptomatic treatment in early stages^3,4^. However, as both the disease and levodopa treatment proceed, patients develop levodopa-induced dyskinesia (LID), characterized by abnormal involuntary movements, and other motor fluctuations, characterized by unpredictable changes in the duration of levodopa’s effects^5,6^. Over years of treatment, these motor complications gradually worsen, constraining treatment and significantly impairing quality of life^7,8^. Clinical research suggests that the progression of motor complications is influenced by three key factors: the extent of neurodegeneration, levodopa dosage, and treatment duration^7,9^. However, disentangling their individual contributions remains challenging, as these factors evolve simultaneously in patients. Despite its clinical significance, the mechanisms driving LID progression and changes in the duration of levodopa’s effect over time are poorly understood, hindering the development of effective therapeutic strategies.

Decades of research in people with PD as well as in animal models of PD have established the striatum as a critical site for LID pathophysiology^10–23^. The principal striatal neurons are GABAergic medium spiny neurons (MSNs), classified according to their dopamine receptor expression: D1 receptor-expressing (D1-MSNs) and D2 receptor-expressing (D2-MSNs)^24^. During movement, D1- and D2-MSNs are coactivated, which has been proposed to be essential for action selection, with D1-MSNs traditionally associated with promoting desired actions and D2-MSNs suppressing competing ones^25–27^. This model suggests that coordinated activity of D1- and D2-MSNs is key for the execution of motor sequences^26,27^. LID can thus be conceptualized as a failure in action selection, where impaired D1- and D2-MSN activity leads to abnormal involuntary movements. Indeed, studies in mouse models of PD have demonstrated that levodopa administration induces opposing changes in the activity of these neurons, by increasing D1-MSN activity and decreasing D2-MSN activity^10,12,13,28,29^. Optogenetic and chemogenetic studies have further established a causal link between D1- and D2-MSN activity and LID expression^12,18,19,30–33^. However, little is known about how levodopa-evoked D1- and D2-MSN responses evolve with repetitive levodopa treatment and how they might contribute to the progression of LID and changes in the duration of levodopa’s effect.

In the healthy brain, dopamine plays a key role in shaping action selection by modulating MSN function through changes in cellular excitability^34,35^ and glutamatergic synaptic plasticity^36–39^. For instance, transient increases and decreases in striatal dopamine in response to action outcomes are thought to reinforce or suppress specific actions by strengthening or weakening corticostriatal synapses^36,40,41^. In PD, the chronic loss of dopamine disrupts this normal process, leading to marked changes in synaptic connectivity in target areas, such as the striatum^14,42^. In late-stage PD with LID, the relationship between dopamine and action outcomes is further disrupted as levodopa leads to a non-physiologic spatiotemporal pattern of striatal dopamine release^43,44^. These sustained increases and decreases in dopamine signaling have been shown to drive a range of cellular adaptations in MSNs, including morphological remodeling, alterations in intrinsic excitability and changes in synaptic connectivity^14,15,22,23,45–48^. Despite these insights, previous research has primarily examined striatal adaptations once LID has already progressed, overlooking the changes that unfold alongside the increase in LID severity. As a result, it is unclear which adaptations drive LID worsening and which are compensatory. We propose that chronic dopamine depletion drives homeostatic but also maladaptive changes in corticostriatal and striatal circuitry, which lead to progressive vulnerability to LID in PD.

Here, we hypothesized that repeated cycles of levodopa-driven dopamine fluctuations promote LID progression by hijacking endogenous mechanisms for striatal synaptic plasticity, leading to changes in D1- and D2-MSN responses to levodopa. To test this hypothesis, we used the 6-hydroxydopamine (6-OHDA) mouse model of PD/LID. By inducing a fixed level of dopamine neuron loss and a constant dose of levodopa (two critical risk factors for LID)^7,9^, this mouse model proved valuable to isolate the effects of repetitive levodopa treatment on LID progression.

Using fiber photometry and single-unit electrophysiology, we tracked changes in D1- and D2-MSN activity, as well as dopamine release, across these critical stages of LID progression. Our findings reveal that repeated levodopa administration is likely to exacerbate LID by amplifying levodopa- induced increases in D1-MSN activity and decreases in D2-MSN activity. Furthermore, using *ex vivo* slice electrophysiology, we demonstrate that LID worsening is associated with cell type-specific potentiation of glutamatergic transmission. Finally, we show that dyskinesia resolves more quickly over the treatment course, which is accompanied by altered D2-MSN activity and reduced D2 receptor sensitivity.

## RESULTS

### Repetitive levodopa treatment worsens LID severity and speeds its onset and offset

To examine how repetitive levodopa administration contributes to LID progression, we used the unilateral 6-OHDA mouse model of PD. After several weeks of recovery, 6-OHDA-lesioned mice were treated with levodopa-benserazide (5 mg/kg-2.5 mg/kg, by daily IP injection) for four weeks (Figure 1A). In response to levodopa, mice increased their overall movement velocity (Figure S1A; see Table S1 for details of the statistical analyses) and developed dyskinesia, which was scored using the abnormal involuntary movement (AIM) scale^49^ at several timepoints across the treatment session (Figure 1A-B). The time course and severity of dyskinesia changed over the course of treatment (Figure 1B), consistent with work by other investigators^17,49^. Most notably, the severity of dyskinesia increased between day 1 and all subsequent days. In addition, the total duration of dyskinesia increased between day 1 and day 4, after which it declined again. To quantify these changes, we focused our analysis on two distinct time windows: **LID onset** (the first 20 minutes after levodopa injection) and **LID offset** (30 to 90 minutes post-injection) (Figure 1B). During the LID onset phase, dyskinesia increased from day 1 to day 4, with no further changes thereafter (Figure 1B-C). In contrast, during the LID offset phase, dyskinesia resolved faster between day 4 and week 4 (Figure 1B, D-E). Levodopa-induced increases in movement velocity also changed over time during both onset and offset phases (Figure S1A-C). During the onset phase, increases in movement velocity were markedly attenuated from day 1 to day 4, which may reflect the fact that severe dyskinesia impairs normal movement (Figure S1A-B). In contrast, during the offset phase, movement velocity gradually increased across weeks (Figure S1C), though the total period of increased velocity was consistent (∼120 minutes). Collectively, these findings indicate that repeated levodopa administration drives two major changes: (1) more rapid onset and higher severity of LID, and (2) more rapid LID offset.

**Figure 1.**
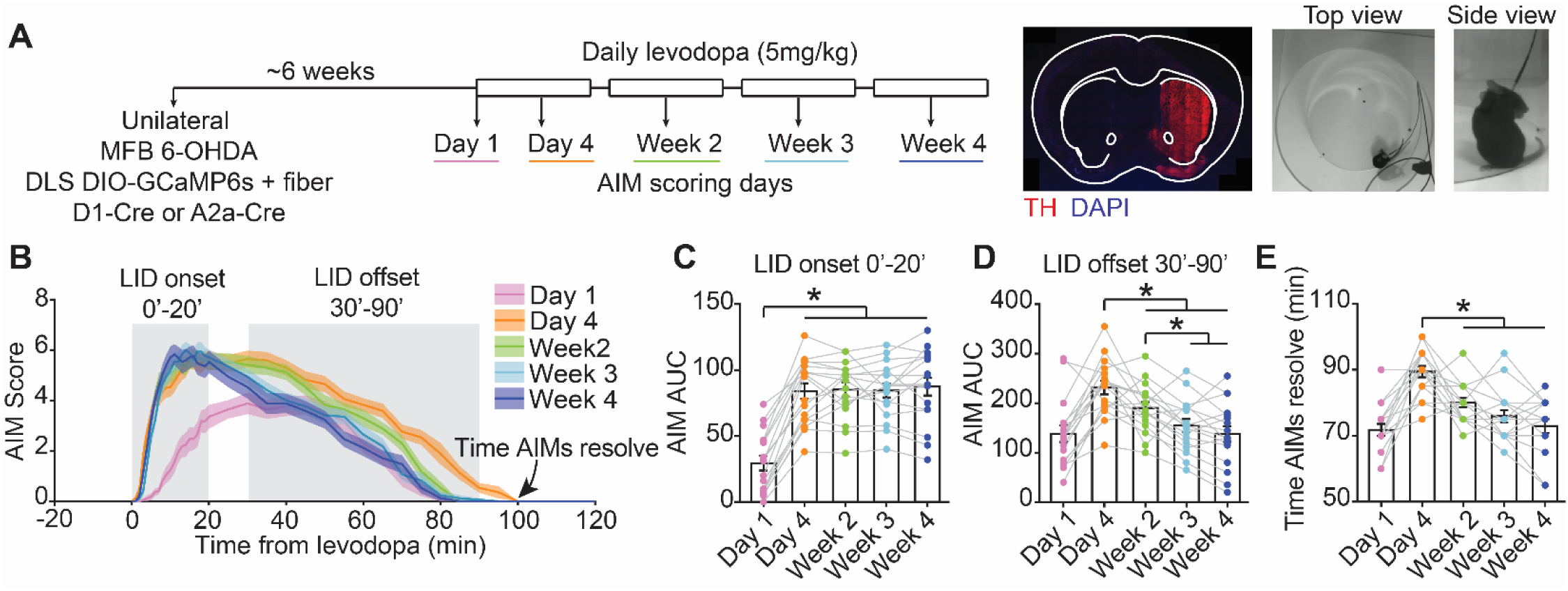
Repetitive levodopa treatment worsens LID severity and speeds its onset and offset. **(A)** Experimental approach. Left: Timeline. Middle: Representative coronal section showing TH loss ipsilateral to 6-OHDA injection. Scale bar: 1mm. Right: Dyskinetic mouse in top and side views. **(B)** Average abnormal involuntary movement (AIM) score after levodopa injection across treatment timepoints. **(C-D)** Area under the curve (AUC) of AIM score during LID onset (C), and offset (D). **(E)** Time to AIM resolution after levodopa injection. [C-E: RM one-way ANOVA, p<0.0001, post-hoc: *p<0.05]. Data shown as mean ± SEM. N=17 mice. See Figure S1.

### Changes in striatal activity parallel alterations in dyskinesia during repetitive levodopa treatment

Neural activity in the striatum has been causally linked to LID^12,18,30^. However, it is unclear how striatal activity evolves with repetitive levodopa administration, and how this activity may contribute to the evolution of dyskinesia over time. To fill this gap, we measured the activity of the two canonical striatal projection neuron types, D1-MSNs and D2-MSNs, over the course of chronic levodopa treatment. We performed cell type-specific fiber photometry with the genetically-encoded calcium indicator GCaMP6s. In 6-OHDA-treated animals, we injected an AAV encoding Cre-dependent GCaMP6s into the dorsolateral striatum (DLS), targeting expression to D1- or D2-MSNs (Figure 2A). Mice were implanted with a 400μm optic fiber in DLS and tested at different timepoints over the course of four weeks of levodopa treatment (Figure 2B). In these recordings, we observed transient elevations of the fluorescence signal, which could be quantified over time and across sessions. Consistent with previous studies using both single-unit electrophysiology and GCaMP calcium imaging^10,28^, levodopa acutely increased D1-MSN activity and decreased D2-MSN activity (Figure 2C).

**Figure 2.**
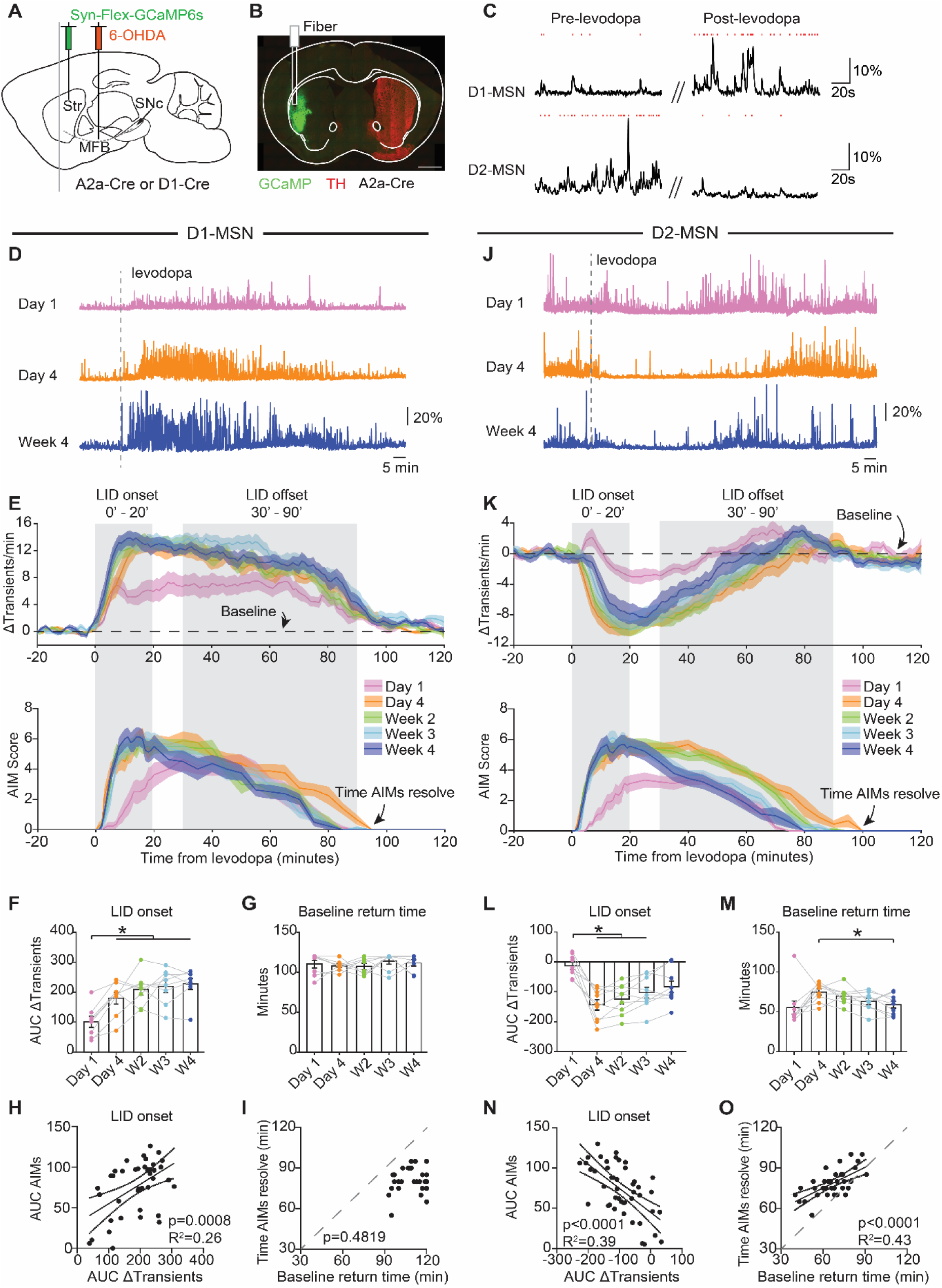
Changes in striatal activity parallel alterations in dyskinesia during repetitive levodopa treatment. **(A)** Schematic of AAV-Flex-GCaMP6s injection into the dorsolateral striatum (Str) and 6-OHDA into the ipsilateral medial forebrain bundle (MFB) of D1-Cre or A2A-Cre mice**. (B)** Representative coronal section showing GCaMP6s expression, TH loss, and fiber track. Scale bar: 1mm. **(C)** Representative short sweeps from GCaMP6s fiber photometry recordings (ΔF/F) from D1- and D2-MSN before and after levodopa, with red ticks marking GCaMP transients. **(D-I)** D1-MSN fiber photometry analysis. **(D)** Representative recordings (ΔF/F) across treatment timepoints (140-minute sessions). **(E)** Top: Change in GCaMP6s transients rate relative to baseline after IP levodopa across treatment timepoints. Bottom: AIM scores for D1-Cre mice. **(F)** Area under the curve (AUC) of Δtransients during LID onset. [RM one-way ANOVA, p=0.0003, post-hoc: *p<0.05]. **(G)** Time for transient frequency to return to baseline. [RM one-way ANOVA, ns]. **(H)** Correlation of Δtransients with AIMs during LID onset. **(I)** Correlation between time of AIMs resolution and the time transient frequency returns to baseline. **(J-O)** D2-MSN fiber photometry analysis. **(J)** Representative recordings (ΔF/F) across treatment timepoints (140-minute sessions). **(K)** Top: Change in GCaMP6s transients rate relative to baseline after IP levodopa across treatment timepoints. Bottom: AIM scores for A2a-Cre mice. **(L)** AUC of Δtransients during LID onset. [RM one-way ANOVA, p=0.0001, post-hoc: *p<0.05]. **(M)** Time for transient frequency to return to baseline. [RM one-way ANOVA, p=0.0047, post-hoc: *p<0.05]. **(N)** Correlation of Δtransients with AIMs during LID onset. **(O)** Correlation between time of AIMs resolution and the time transient frequency returns to baseline. N=8 D1-Cre and N=9 A2a-Cre. Each dot is a session of one mouse. Data shown as mean ± SEM (E-G; K-M). See Figures S2, S3 & S4.

We next compared the response of each striatal subpopulation to levodopa across 1 month of repeated administration. Given the close linkage of increased D1-MSN activity to LID^10,12,13,19,28–30,50,51^, we hypothesized that increases in D1-MSN activity across sessions might parallel the worsening of LID over time. We compared D1-MSN responses to levodopa at day 1, day 4, and weeks 2-4. We measured these responses by comparing the fluorescence transient frequency (Figure 2D-E). Importantly, baseline transient rates (pre-levodopa administration) were unchanged by repetitive treatment (Figure S2A). In the onset phase, we observed parallel increases in dyskinesia severity and D1-MSN activity between day 1 and day 4 onward (Figure 2E-F). In the offset phase, we observed a more rapid offset and faster resolution of LID between day 4 and week 4, but this behavioral change was not paralleled by any change in D1-MSN activity (Figure S2B; Figure 2E). Moreover, the time required for D1-MSN activity to return to pre-levodopa baseline levels remained constant across treatment timepoints (Figure 2G). To further examine the link between D1-MSN activity and AIM scores during LID onset and offset phases, we correlated the area under the curve (AUC) of the AIM score with the change in transient frequency. Remarkably, D1-MSN activity showed a robust correlation with AIM scores during the LID onset, but not during the offset phase (Figure 2H; Figure S2C). In line with this result, the time required for D1-MSN activity to return to pre-levodopa baseline levels did not correlate with the time that AIMs resolved (Figure 2I). Together, these findings indicate D1-MSN activity parallels the worsening of dyskinesia severity, but may not explain why LID resolves more rapidly at later timepoints in treatment.

D2-MSNs have also been closely linked to LID, as chemogenetic or optogenetic activation attenuates LID severity^18,30^. We hypothesized that changes in LID severity and kinetics might also be mediated by changes in D2-MSN responses to levodopa over time. While D2-MSN activity was reduced by levodopa at all time points, this suppression was more pronounced by day 4 of treatment (Figure 2J-L). This suppression was prominent during the LID onset phase, paralleling the increase in LID severity from day 1 to day 4 (Figure 2K-L), with no changes in baseline D2-MSN GCaMP transient frequency (Figure S2D). Moreover, the recovery of D2-MSN activity was more rapid during the offset phase from day 4 to week 4 (Figure 2M; Figure S2E), which paralleled the more rapid resolution of AIMs (Figure 2K). Indeed, D2-MSN activity was strongly correlated with AIM scores for both LID onset and offset periods (Figure 2N; Figure S2F), and the time required for D2-MSN activity to return to pre-levodopa baseline levels showed robust correlations with the time required for AIMs to resolve (Figure 2O). Importantly, we obtained identical results when using ΔF/F amplitudes, instead of transient frequency, as a proxy of neural activity, both for D1- and D2-MSNs (Figure S3). These results demonstrate that alterations in D2-MSN responses to levodopa parallel changes in the onset and offset of LID over the course of treatment.

To examine the relationship between MSN activity and behavior more systematically, we compared calcium transient frequency across three behavioral states throughout the levodopa session: pre-levodopa hypokinesia, prokinetic periods without dyskinesia, and dyskinetic periods (Figure S4A). We found that during hypokinesia, D2-MSN activity predominated; during prokinetic periods without dyskinesia, D1- and D2-MSN activity was relatively balanced; and during dyskinesia, activity was biased towards D1-MSNs. Moreover, the difference between average D1- and D2-MSN GCaMP transient frequencies strongly correlated with dyskinesia severity across AIM scores (Figure S4B-C). Together, these results show that the relative activity of D1- and D2-MSN is closely associated with behavioral state and dyskinesia severity.

### Repetitive levodopa potentiates the responses of single striatal neurons, without changes in dopamine release

GCaMP fiber photometry allows measurement of cell-type specific neural activity in a stable population of neurons over the course of many sessions. However, it reflects population, rather than single-cell activity. Changes in bulk D1- and D2-MSN GCaMP signal after repetitive levodopa may result from alterations in ensemble size, amplitude of single-cell responses, or both. To distinguish these possibilities, we performed two new sets of experiments. First, we assessed single-cell activation with a surrogate marker, the immediate early genes c-Fos and ΔFosB. These studies allowed us to approximate the number of cells recruited by levodopa at different timepoints, and determine whether this number changes over time. Second, we performed single-unit electrophysiology to determine whether repetitive treatment changes the firing rates of single striatal neurons. Administration of levodopa has been shown to induce immediate early gene expression in the striatum^50,52–54^. Specifically, c-Fos is increased in a subpopulation of striatal neurons, composed primarily of D1-MSNs, which are causally linked to LID^12,13^. To determine if a larger ensemble of D1-MSNs is recruited over the course of treatment, we sacrificed 6-OHDA-treated Drd1-tdTomato mice 2 hours after levodopa injection on day 1, day 4 or week 4 of treatment, and immunostained for c-Fos or ΔFosB. Consistent with previous studies^12^, the majority of c-Fos+ or ΔFosB cells were D1-MSNs (84-91% for c-Fos and 70-75% for ΔFosB), as measured by colocalization with tdTomato. We found that the density of c-Fos+ or ΔFosB and colocalized c-Fos/tdTomato+ or ΔFosB/tdTomato+ cells in the dorsolateral striatum did not differ between mice receiving varying durations of levodopa treatment (Figure 3A-F). These results suggest that a similar number of D1-MSNs are recruited by each levodopa administration, regardless of treatment duration.

**Figure 3.**
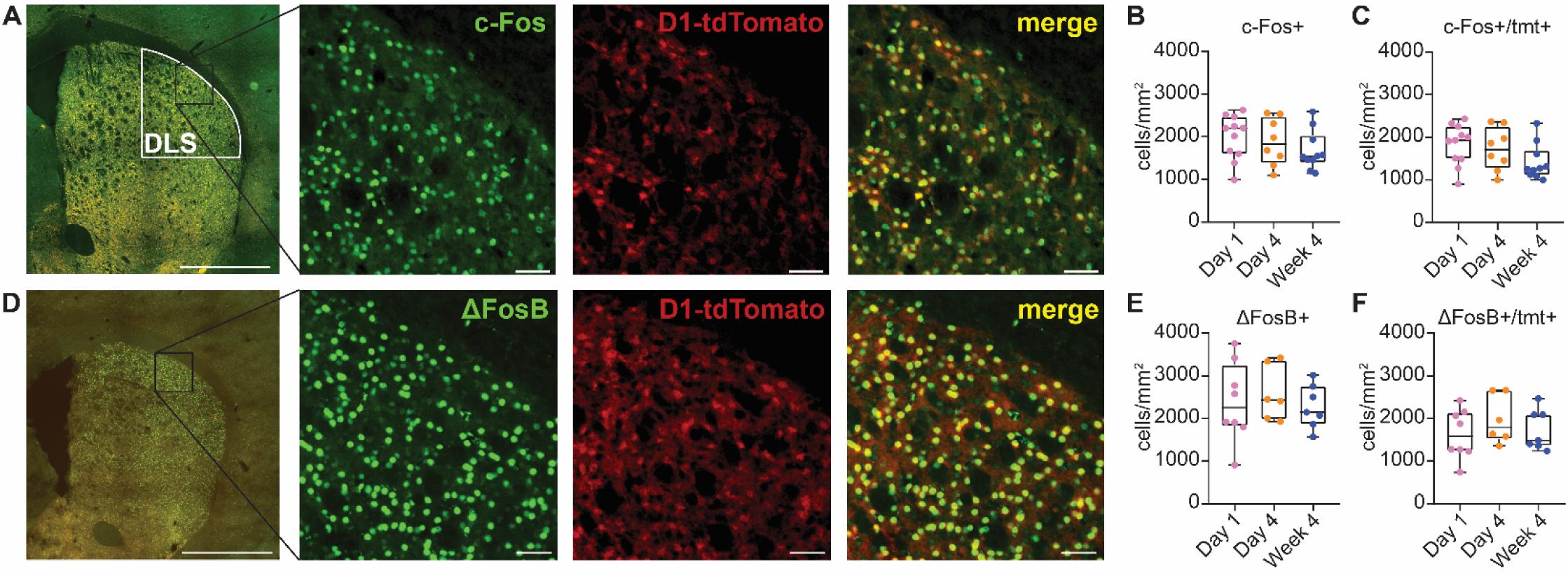
Repetitive levodopa treatment recruits a similar number of D1-MSNs over time. **(A)** Representative coronal section showing D1-tdTomato and c-Fos immunofluorescence in the dorsolateral striatum (DLS). Scale bar: 1mm. Inset: Magnified c-Fos, D1-tdTomato, and merged images. Scale bar: 50µm. **(B-C)** Density of c-Fos+ cells (B) and colocalized c-Fos+/D1+ cells (C) across timepoints. [Kruskal-Wallis, ns]. Each dot represents one mouse (N=8-11). **(D)** Representative coronal section showing D1-tdTomato and ΔFosB immunofluorescence in the dorsolateral striatum. Scale bar: 1mm. Inset: Magnified ΔFosB, D1-tdTomato, and merged images. Scale bar: 50µm. **(E-F)** Density of ΔFosB+ cells (E) and colocalized ΔFosB+/D1+ cells (F) across timepoints. [Kruskal-Wallis, ns]. Each dot represents one mouse (N=6-8). N=animals. Data shown as median and range (B,C,E,F).

To determine if single-cell firing responses to levodopa change over repetitive administration, we used *in vivo* single-unit electrophysiology in freely-moving parkinsonian mice. 6-OHDA-treated mice were implanted with an electrophysiology recording device in the DLS (Figure 4A). We performed single-unit recordings during levodopa treatment sessions on day 1, days 3-5 and week 4 of levodopa treatment. We identified and excluded putative interneurons based on spike waveform^10,55–57^. In line with previous findings^10^, we observed a variety of responses to levodopa across striatal neurons, including increased firing, decreased firing and no response. Based on previous optically labeled recordings in the same mouse model^10^, units that increased or decreased their firing in response to levodopa were classified as putative D1- or D2-MSNs, respectively (see Methods). No response units, which represented 9% of all units, were not analyzed further. Over time, the onset phase showed greater differences in dyskinesia severity and striatal GCaMP signal over time, so we focused on this phase in our analysis of single-unit firing responses. As the number of levodopa-activated D1-MSNs appeared to be stable across 4 weeks of treatment, we hypothesized that the levodopa-evoked firing of individual neurons would increase, producing the potentiation of D1-MSN GCaMP responses across treatment. Indeed, putative D1-MSNs showed greater levodopa-evoked increases in firing at the single-unit level (Figure 4B-E). Putative D2-MSNs also showed lower absolute firing rates during LID onset after repetitive levodopa treatment (Figure S5A-C), although the normalized firing rate relative to baseline was not significantly different over time (Figure S5D-E). Moreover, paralleling our GCaMP data, relative D1-/D2-MSN activity, as measured by firing rates, was strongly correlated with dyskinesia severity (Figure S4D-E).

**Figure 4.**
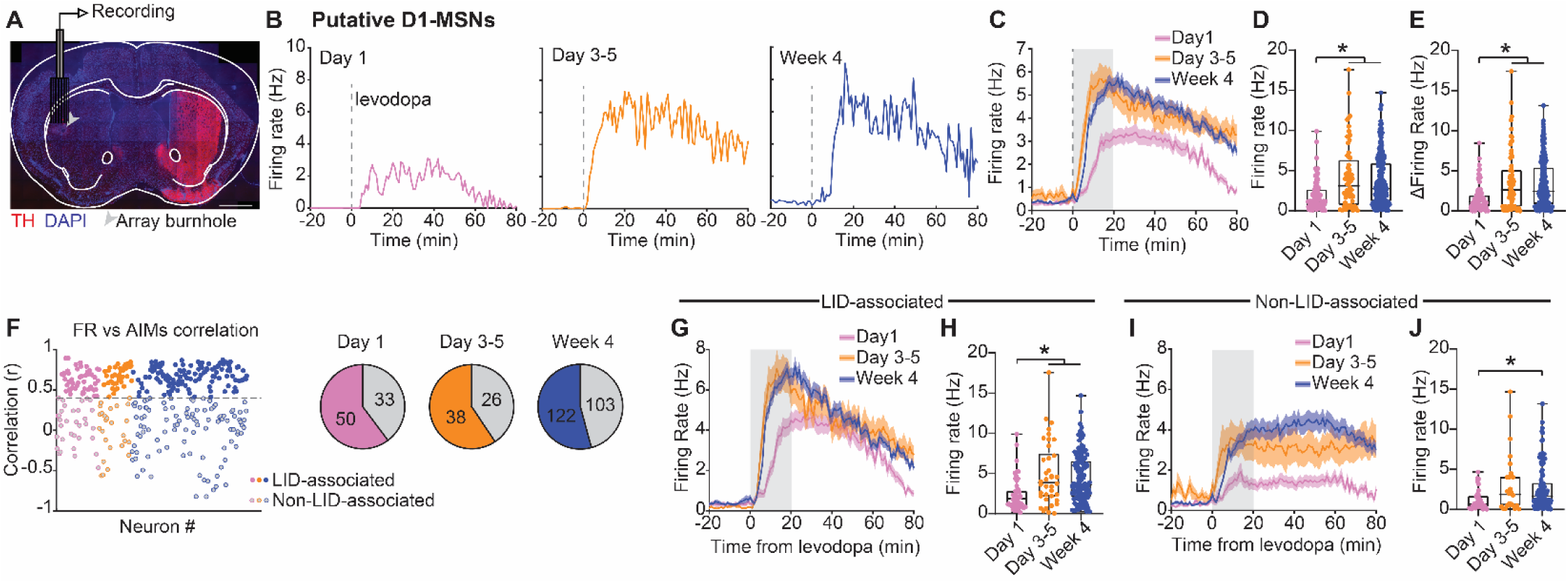
Repetitive levodopa treatments potentiate single D1-MSN neuron responses. **(A)** Schematic of multielectrode array implant in DLS and postmortem histology showing recording site with electrolytic lesion. **(B-C)** Representative single units (B) and pooled firing rates (C) of putative D1-MSN on days 1, 3-5 and week 4 aligned to levodopa injection. **(D-E)** Average firing rate (D) and change relative to baseline (E) during LID onset (gray bar in C). [Kruskal-Wallis, p<0.0001, post-hoc: *p<0.001]. Day 1: n=83, N=11; Day 3-5: n=64, N=11; Week 4: n=225, N=15. **(F)** *Left:* Distribution of correlations between putative D1-MSN firing rates and AIM scores across treatment days. Filled dots indicate units classified as LID-associated. *Right:* Proportion of putative LID-associated (colored) and non-LID-associated (gray) D1-MSNs across treatment days. **(G-H)** Firing rates of LID-associated D1-MSNs in response to levodopa injections (G) and average firing rates during LID onset (H). [Kruskal-Wallis: p<0.0001, post-hoc: *p<0.001]. Day 1: n=50, N=11; Day 3-5: n=38, N=11; Week 4: n=122, N=15. **(I-J)** Firing rates of non-LID-associated D1-MSNs in response to levodopa injections (I) and average firing rates during LID onset (J) [Kruskal-Wallis: p=0.0091, post-hoc: *p=0.0069]. Day 1: n=33, N=11; Day 3-5: n=26, N=11; Week 4: n=103, N=15. n=units; N=animals. Data shown as mean ± SEM (C,G,I) and as median and range (D,E,H,J). See Figures S4, S5 & S6.

In previous work, we found that the firing rate of a subset of D1-MSNs was highly correlated with LID severity (“LID-associated” cells) and that these neurons exhibited abnormally high levodopa-evoked firing rates^10,13^. To determine whether the changes in firing rates after repetitive treatment reflect an expansion in the proportion of LID-associated cells, or instead a broader increase in firing across D1-MSNs, we classified both D1-MSNs and D2-MSNs as LID-associated and non-LID-associated based on their correlation with AIMs (see Methods). The proportion (and distribution of correlation strengths) of LID-associated D1-MSNs did not significantly increase with repetitive treatment (Figure 4F). Though LID-associated and non-LID-associated D1-MSNs have distinct firing profiles across individual levodopa treatment sessions, both populations increase their firing rates over the course of treatment (Figure 4G-J). Notably, among D2-MSNs, only LID-associated neurons exhibited changes in firing rate following repetitive treatment (Figure S5F-J). Finally, consistent with our fiber photometry recordings, both LID-associated D1- and D2-MSNs were correlated with AIMs during the onset phase, whereas during the offset phase only D2-MSNs remained significantly correlated (Figure S6).

Taken together, these findings suggest that repetitive levodopa administration primarily alters D1- MSN responses through changes in the magnitude of single-cell activity rather than through recruitment of additional LID-associated neurons.

We next sought to investigate the cellular mechanisms driving changes in D1- and D2-MSN responses to levodopa over repetitive treatment. Given that bidirectional levodopa responses of D1- and D2-MSNs potentiate in parallel between day 1 and day 4, one plausible mechanism is alterations in dopamine signaling. To determine whether levodopa-evoked dopamine release changes over this period, we performed fiber photometry with the fluorescent dopamine sensor GRAB-DA2h^58^ in wild type 6-OHDA-treated mice (Figure 5A-B). We recorded dopamine release, as measured by GRAB-DA2h, in response to levodopa across four weeks of treatment. As expected, each levodopa administration increased dopamine release, with a time course that roughly matched the duration of increased movement and dyskinesia (Figure 5C). These responses were consistent with slow and sustained increases in dopamine, rather than with fast transient elevations. Most notably, the amplitude and timecourse of levodopa-evoked dopamine release was consistent over four weeks of treatment (Figure 5D-E). These results suggest that changes in dopamine release are unlikely to explain the potentiation of MSN responses over the course of treatment.

**Figure 5.**
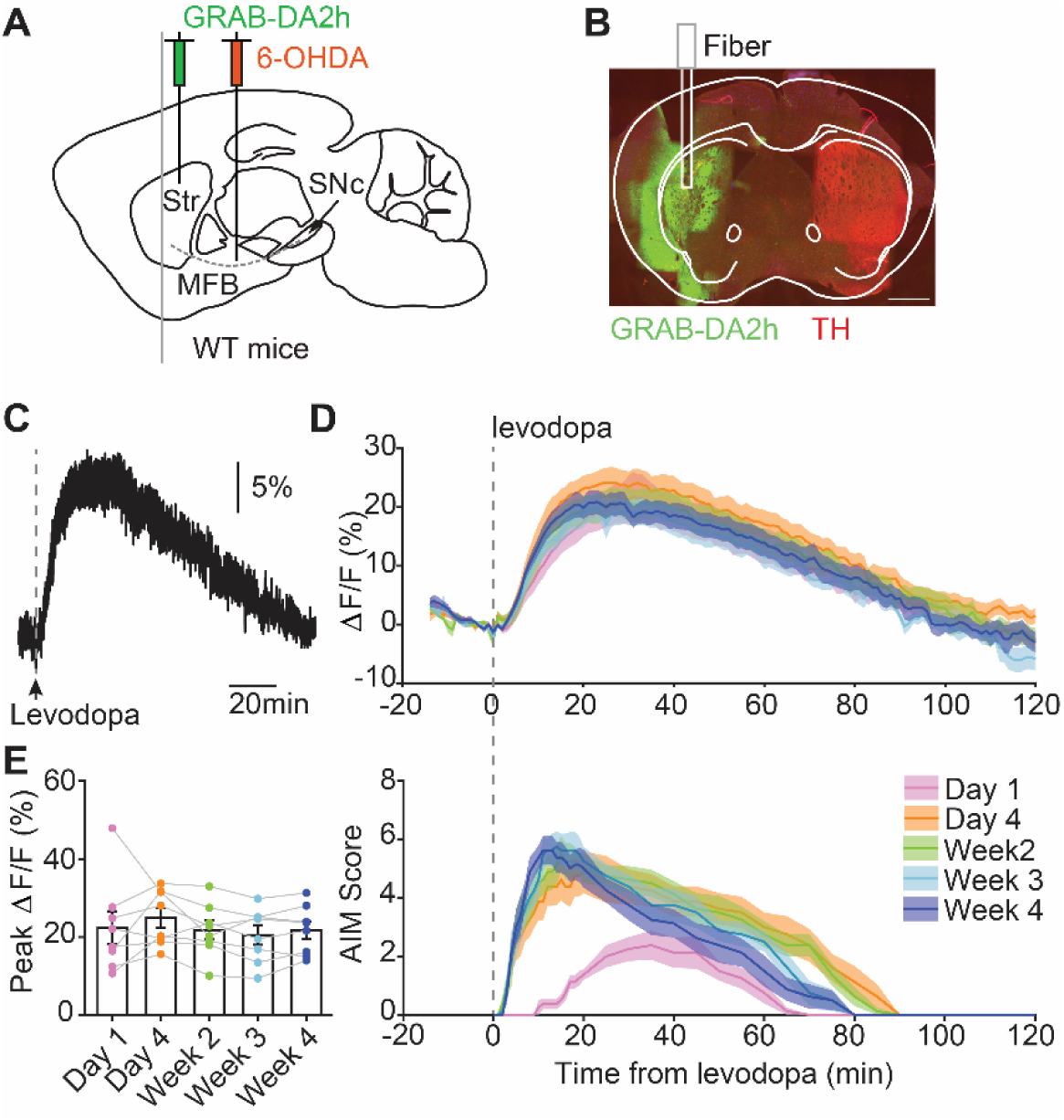
Repetitive levodopa does not alter striatal dopamine release. **(A)** Schematic of AAV-GRAB-DA2h injection and fiber implant in the dorsolateral striatum, and 6-OHDA injection into the MFB of WT mice. **(B)** Representative coronal section showing GRAB-DA2h expression and TH loss. **(C)** Representative GRAB-DA2h fiber photometry signal in response to levodopa. **(D)** Average normalized GRAB-DA2h (ΔF/F) responses to levodopa across treatment timepoints (top), aligned to AIM scores (bottom). **(E)** Peak normalized GRAB-DA2h ΔF/F responses to levodopa across timepoints. [RM one-way ANOVA, ns]. N=8 mice. Each dot is a session of one mouse.

### Repetitive levodopa administration increases excitatory transmission in D1-MSNs and reduces D2R sensitivity in D2-MSN

Though levodopa-evoked striatal dopamine release was consistent across treatment sessions, these repeated elevations may lead to changes in the properties of downstream circuit elements. One of the most well-known mechanisms through which dopamine can induce long-lasting alterations in striatal function is by controlling glutamatergic synaptic plasticity^36–38^. Transient increases in striatal dopamine in response to positive action outcomes drive the induction of corticostriatal synaptic plasticity^36,37,59^, thereby increasing the probability of repeating that action in the future^37,59^. In the dopamine-depleted striatum, the relationship between dopamine and action outcomes is disrupted, with exceptionally low striatal dopamine in the untreated state, and with levodopa treatment, dopamine increases for minutes to hours. These alterations drive aberrant plasticity at striatal glutamatergic synapses after weeks of levodopa treatment^15,48^. However, it is unclear how striatal glutamatergic synapses are shaped over the course of levodopa treatment and how they relate to the progression of LID. To fill this gap, we performed *ex vivo* slice recordings in Drd1-tdTomato or D2-GFP 6-OHDA-treated mice, to allow targeting of D1- and D2-MSNs. We performed recordings in three treatment groups: treatment-naïve animals (mimicking baseline conditions on day 1 of levodopa), animals treated with levodopa for 3 days or 4 weeks (mimicking day 4 and week 4). Brain slices were prepared 24 hours after the last dose of levodopa, corresponding to the “OFF” levodopa state. This timing avoids acute effects of dopamine receptor activation during the “ON” state, thereby isolating chronic effects of repetitive levodopa exposure. To assess glutamatergic synaptic transmission onto D1- and D2-MSNs, we measured miniature excitatory postsynaptic currents (mEPSCs) in the presence of tetrodotoxin and picrotoxin (Figure 6A). Compared to the levodopa-naïve condition, we found an increase in the frequency of mEPSCs in D1-MSNs at both day 4 and week 4 of levodopa treatment (Figure 6B). There were small changes in mEPSC amplitudes, only evident between levodopa-naïve and week 4 groups (Figure 6C). In contrast, mEPSC frequency and amplitude in D2-MSNs were unchanged after repetitive levodopa (Figure 6D-F).

**Figure 6.**
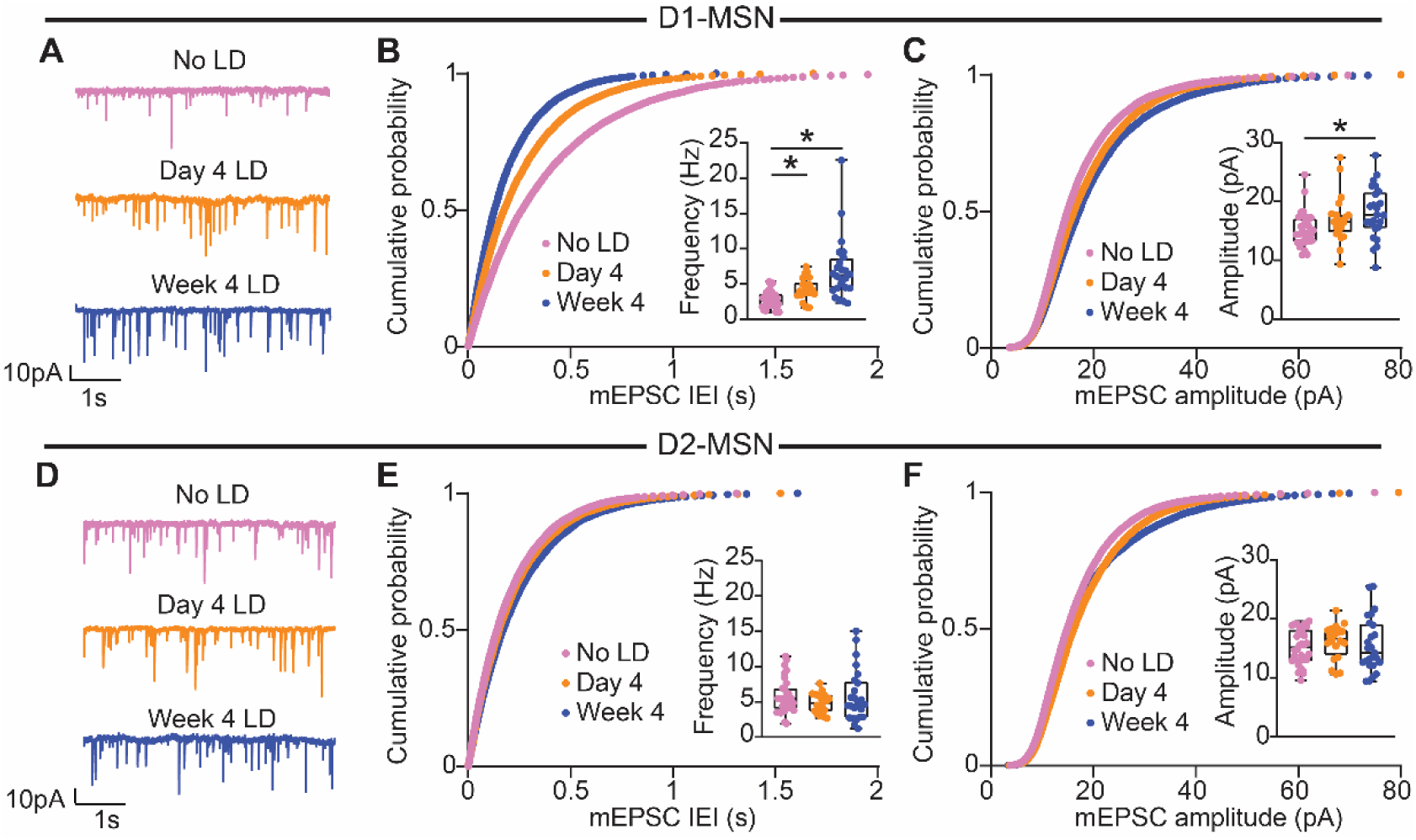
Repetitive levodopa treatments drive increased excitatory transmission onto D1-MSNs. **(A)** Representative voltage-clamp recordings of miniature excitatory postsynaptic currents (mEPSC) in D1-MSNs across treatment timepoints. **(B)** Cumulative probability distribution of mEPSC Inter-Event-Intervals (IEI) in D1-MSNs. Inset: average mEPSC frequency. [Kruskal-Wallis, p<0.0001, post-hoc comparisons *p<0.05]. **(C)** Cumulative probability distribution of mEPSC amplitudes in D1-MSNs. Inset: average mEPSC amplitude. [Kruskal-Wallis, p=0.0158, post-hoc comparisons *p=0.0150]. No LD: n=30, N=11; Day 4 LD: n=22, N=6; Week 4 LD: n=24, N=6. **(D)** Representative voltage-clamp recordings of mEPSC in D2-MSNs across treatment timepoints. **(E)** Cumulative probability distribution of mEPSC Inter-Event-Intervals (IEI) in D2-MSNs. Inset: average mEPSC frequency. [Kruskal-Wallis, ns]. **(F)** Cumulative probability distribution of mEPSC amplitudes in D2-MSN. Inset: average mEPSC amplitude. [Kruskal-Wallis, ns]. No LD: n=32, N=10; Day 4 LD: n=20, N=6; Week 4 LD: n=25, N=6. n=cells; N=mice. Data shown as mean for probability plots, and median and range for insets. See Figure S7.

To determine whether this increase in glutamatergic drive onto D1-MSN is also present in the levodopa “ON” state more closely modeling periods of LID, we prepared slices of 6-OHDA-treated Drd1-tdTomato mice 30 minutes after levodopa injection and assessed excitatory synaptic transmission during this simulated “ON” state^22^. Consistent with the “OFF” state results, both the frequency and amplitude of mEPSCs in D1-MSNs were increased at day 4 and week 4 of levodopa treatment (Figure S7). This increased D1-MSN mEPSC frequency and amplitude paralleled the potentiation in D1-MSN responses to levodopa and LID severity from day 1 to day 4, making glutamatergic synaptic plasticity a strong candidate mechanism underlying LID progression.

Aside from controlling synaptic plasticity, dopamine can also influence D1- and D2-MSN activity by modulating cellular excitability through regulation of multiple ion channels^34,60,61^. Previous studies have shown that dopamine neuron degeneration and subsequent levodopa therapy can induce chronic alterations in MSN excitability^14,45^. To directly test whether D1- and/or D2-MSN intrinsic excitability is altered after four days or four weeks of levodopa treatment, we again made *ex vivo* whole-cell patch clamp recordings from identified D1- or D2-MSNs in 6-OHDA-treated mice. In current-clamp recordings, we gave a series of current injections to evoke spiking (Figure 7A), and plotted the number of action potentials as a function of input current, as well as the rheobase. We found that D1-MSN excitability was unchanged after four days of levodopa administration, when the severity of LID has already peaked (Figure 7B-C). However, we found a marked reduction in excitability after four weeks of treatment, manifested as a reduced number of action potentials in response to depolarizing current steps and an increase in rheobase (Figure 7A-C). As we did with synaptic physiology, we repeated these experiments in a simulated “ON” state^22^, and found that these differences in D1-MSN excitability between timepoints persisted (Figure S8). In contrast, D2-MSNs did not show alterations in intrinsic excitability after four days or four weeks of treatment (Figure 7D-F). These results suggest changes in intrinsic excitability are unlikely to explain changes in D1-/D2-MSN activity or LID severity after repetitive treatment.

**Figure 7.**
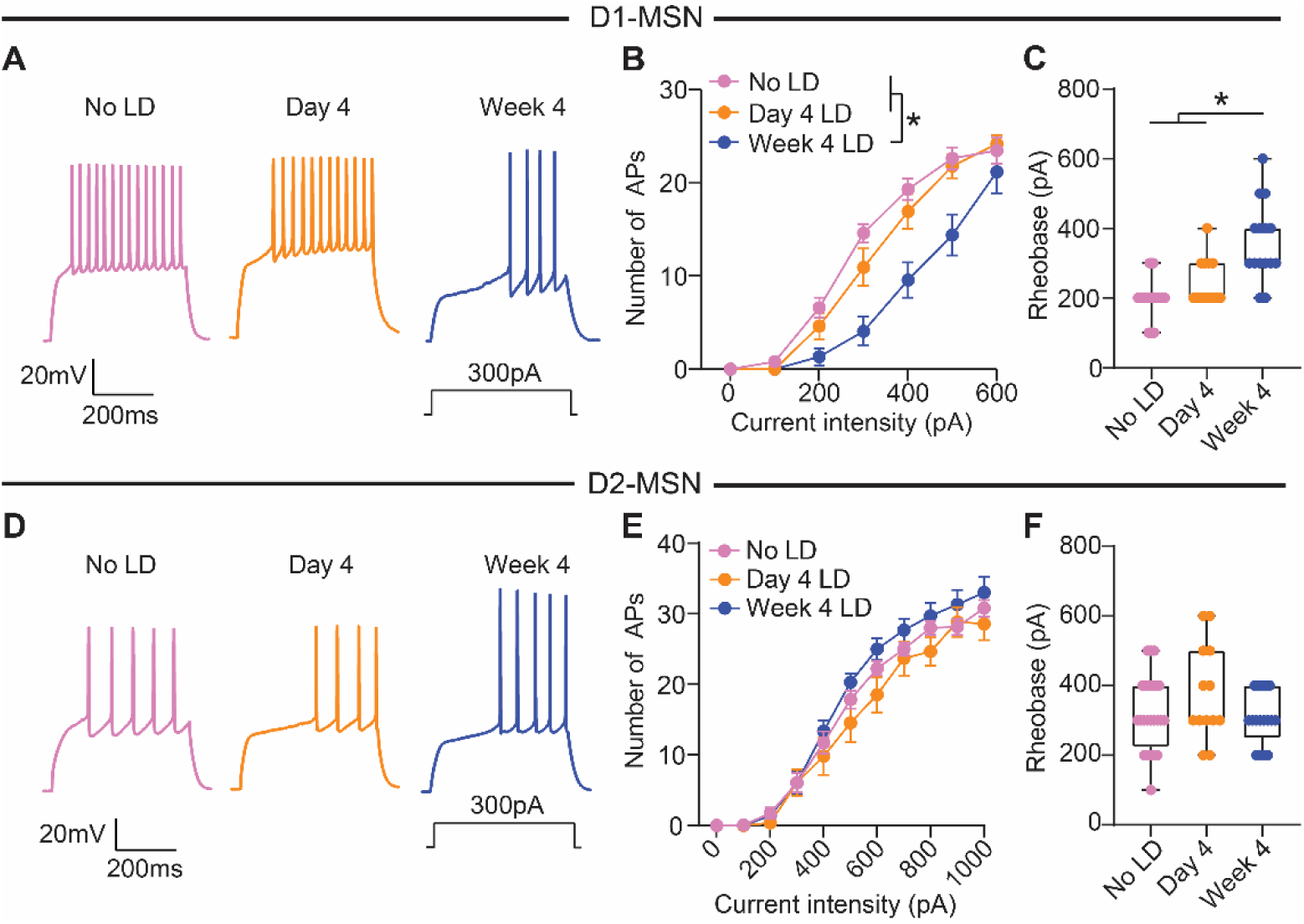
D1-MSN intrinsic excitability is reduced after repetitive levodopa treatment. **(A)** Voltage responses to current injection in D1-MSNs across treatment timepoints. **(B)** Number of action potentials (APs) in response to depolarizing current steps. [Two-way ANOVA, treatment p=0.0001, interaction p<0.0001, post-hoc for 300, 400 and 500pA *p<0.05]. **(C)** D1-MSN rheobase across treatment duration. [Kruskal-Wallis, p<0.0001, post-hoc *p<0.05]. No LD: n=22, N=9; Day 4: n=13, N=6; Week 4: n=15, N=6. **(D)** Voltage responses to current injection in D2-MSN across treatment timepoints. **(E)** Number of APs in response to depolarizing current steps. [Two-way ANOVA, ns]. **(F)** D2-MSN rheobase across treatment duration. [Kruskal-Wallis, ns]. No LD: n=24, N=11; Day 4: n=13, N=6; Week 4: n=17, N=5. n=cells; N=animals. Data shown as mean ± SEM (B,E) and as median and range (C,F). n=cells; N=mice. Each dot represents one cell. See Figures S8 & S9.

We next investigated candidate ionic conductances underlying reduced D1-MSN excitability after 4 weeks of treatment. Lower excitability often correlates with increased potassium conductances, including Kv1.2-containing channels^62^ and calcium-activated small-conductance (SK) and big-conductance (BK) potassium channels that mediate the afterhyperpolarization (AHP)^63–65^. Although conclusive identification of membrane currents requires voltage-clamp measurements, examination of membrane properties can suggest candidate currents^34^. While D1-MSN action potential threshold was unchanged, AHP amplitude was increased and spike half-width was decreased after four weeks of treatment (Figure S9A-D), consistent with BK/SK upregulation. Resting membrane potential and responses to hyperpolarizing current, mainly determined by inwardly rectifying Kir2 channels^66^, were also unchanged (Figure S9E-F). However, subthreshold responses to depolarizing current injections were attenuated after four weeks of treatment (Figure S9F), consistent with possible upregulation of Kv1.2-containing channels that mediate A-type potassium currents, I ^61^. D2-MSN membrane properties remained unchanged (Figure S9G-L), in line with unchanged intrinsic excitability. These results suggest potassium currents as one potential mechanism underlying decreased D1-MSN excitability over levodopa treatment.

While we identified a candidate mechanism for potentiated D1-MSN responses, we found no changes in D2-MSN intrinsic or synaptic properties that might explain their enhanced suppression on day 4, and the progressively faster recovery from day 4 to week 4. One alternative explanation is alteration in dopamine signaling through dopamine D2 receptors (D2R). To measure D2R signaling with an electrophysiological readout, we virally overexpressed an effector of D2R signaling, the G-protein-activated inwardly rectifying potassium channel (GIRK2)^67,68^ together with tdTomato in D2-GFP mice (Figure S10A). Mice were either left untreated (no LD), or received levodopa for 3 days or 4 weeks. We performed whole-cell voltage-clamp recordings (Vh=-60mV) from GFP+/tdTomato+ cells 24 hours after the last dose. Bath-applied dopamine induced D2R-mediated outward GIRK currents (Figure S10B-C). Responses to 1μM dopamine (near the EC50 in levodopa-naïve 6-OHDA-treated mice) were normalized to those at saturating concentration (100μM) to account for intercellular variability in GIRK expression (Figure S10C-F). We found that the sensitivity of D2R to dopamine was unchanged on day 4, as evidenced by a stable ratio of GIRK currents relative to the levodopa-naïve condition (Figure S10F). Thus, changes in D2R sensitivity cannot explain the enhanced suppression of D2-MSNs observed already at day 4 of treatment. However, after 4 weeks, a reduced GIRK current ratio suggest a decrease in D2R sensitivity (Figure S10F), which paralleled the faster D2-MSN recovery and AIMs resolution.

## DISCUSSION

In this study, we combined behavior with *in vivo* and *ex vivo* techniques to understand the physiological correlates of progressive LID over the course of a month of levodopa therapy. We found that repetitive levodopa administration drives two major changes in behavior: (1) worsening severity and more rapid onset of LID, and (2) accelerated resolution of LID. Moreover, we revealed that these alterations are paralleled by distinct long-lasting alterations in striatal function: (1) worsening dyskinesia severity and more rapid onset correlated with potentiation of D1-MSN and D2-MSN responses to levodopa and with enhanced glutamatergic synaptic inputs onto D1-MSNs; (2) more rapid LID offset is linked to more rapid recovery of D2-MSN activity and reduced D2R sensitivity (Figure 8).

**Figure 8.**
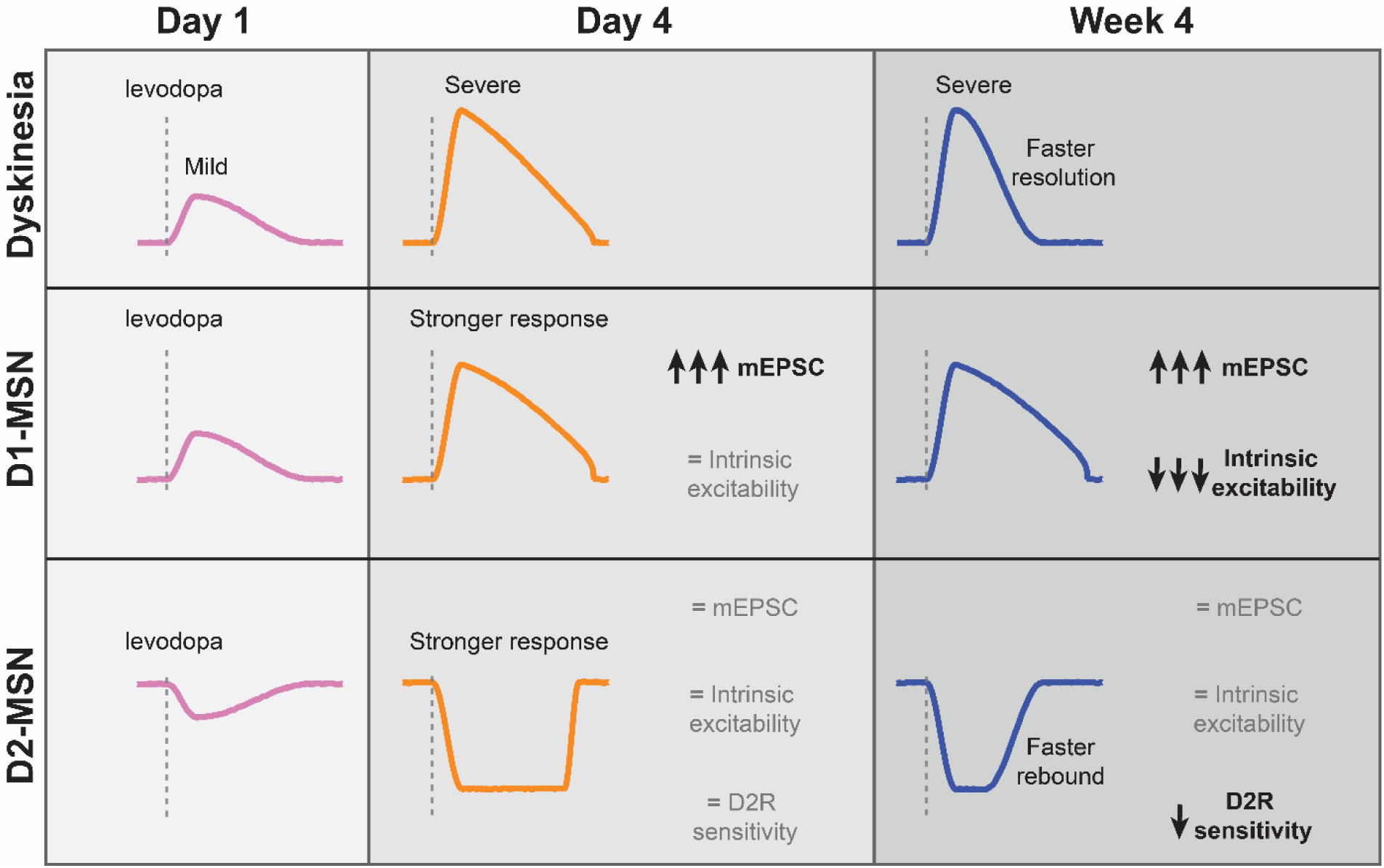
Summary of D1-MSN and D2-MSN adaptations following repetitive levodopa treatment.

People with PD gradually develop LID, which then worsens over years of levodopa treatment. Clinical studies suggest that advancing neurodegeneration, treatment duration and levodopa dosage are key risk factors for LID development and worsening^7,9^. However, disentangling their individual contributions remains challenging, as these factors evolve simultaneously. Using the 6-OHDA mouse model, we isolated the effects of repetitive levodopa treatment, as the extent of neurodegeneration and dosage remained fixed. This model differs from human PD in its rapid development of LID, likely due to near-complete dopamine depletion and the levodopa dose used, which may accelerate the underlying plasticity mechanisms. Nevertheless, repeated levodopa exposure (at least 2-4 days) is required to maximize LID severity in this model, and changes in LID duration continues to evolve over weeks. Therefore, while the temporal dynamics differ, the cellular and synaptic mechanisms engaged in our model may reflect those operating more slowly in patients.

Our results demonstrate that repetitive levodopa worsens LID, resulting in more rapid onset and severe dyskinesia. Based on prior studies linking D1-MSNs activity causally to LID^12,19,69^, we hypothesized this worsening might be accompanied by increased levodopa-evoked D1-MSN activity. Indeed, both GCaMP fiber photometry and single-unit electrophysiology revealed progressively enhanced D1-MSN responses, at both the population and single-cell level. Though immediate early gene expression may not fully capture the ensemble of D1-MSNs activated by levodopa, we did not see a change in the number of c-Fos- or ΔFosB-labeled D1-MSNs in dyskinesia over the course of treatment, in line with previous observations^70^. Our findings suggest that potentiation of D1-MSN firing rate responses to levodopa could contribute to the worsening of LID over time.

The potentiation of D1-MSN responses observed *in vivo* could reflect an increase in dopamine signaling, intrinsic excitability, and/or synaptic plasticity. GRAB-DA2h fiber photometry showed no detectable changes in dopamine release, although this approach may not be able to resolve small differences in dopamine release, or account for spatial heterogeneity. Alternative techniques, such as fast-scan cyclic voltammetry^71^, or spatially resolved imaging^72,73^ might better capture changes in dopamine release dynamics. As dopamine increases the excitability of D1-MSNs via D1 receptors^22,34,74^, the increase in D1-MSN activity may be mediated by enhanced postsynaptic responses.

Increased intrinsic excitability could explain the higher levodopa-evoked D1-MSN firing rates, but we did not find evidence for increased excitability. Instead, we found heightened excitatory synaptic input that paralleled LID progression. Strikingly, this increase in excitatory inputs did not produce higher D1-MSNs baseline firing or calcium activity in the absence of levodopa. Although synaptic inputs may be stronger, they may fail to drive spiking when presynaptic activity or postsynaptic excitability is low, as is typical for MSNs. Thus, these synaptic changes may become functionally relevant only after levodopa administration, when dopamine enhances D1-MSN excitability and responsiveness to excitatory input. Indeed, previous observations indicate that despite differences in anatomical and intrinsic properties between D1- and D2-MSNs in healthy mice^75^, both populations show similarly low baseline firing rates *in vivo*^10^, suggesting that differences in *ex vivo* measures may only translate into altered in *in vivo* firing under specific network conditions. Previous studies found evidence for increased synaptic strength in animals with LID^14,15,48^, but did not consider the effect of time as we did here. Repeated and prolonged elevation of striatal dopamine may hijack existing mechanisms for striatal synaptic plasticity, promoting aberrant potentiation of excitatory synapses onto D1-MSNs^15,48^. Dopamine-dependent potentiation of corticostriatal synapses onto D1-MSNs is key for reinforcing actions associated with positive outcomes^37,76^. In LID, dopamine becomes uncoupled from natural action-outcome associations. Therefore, rather than selectively enhancing plasticity in action-relevant D1-MSNs, sustained dopamine elevations may promote aberrant plasticity preferentially impacting a subset of D1-MSNs that may be intrinsically more vulnerable (i.e., the “LID-associated cells” exhibiting abnormally high firing rates). Elucidating the mechanisms underlying this maladaptive plasticity will be critical for developing effective antidyskinetic therapies and should be a priority for future research.

Although D1-MSNs play a crucial role in LID, D2-MSNs have also been implicated^18,22,23,30^. In parallel with enhanced D1-MSN responses, we found a reciprocal change in D2-MSN activity: stronger suppression by levodopa over time. We observed this change using GCaMP fiber photometry, while with single-unit electrophysiology it was harder to resolve a difference in firing rates relative to baseline. This discrepancy may reflect technical limitations. First, our classification of “putative D1- or D2-MSNs” based on firing rate changes in response to levodopa likely biases the detection rate toward D1-MSNs, which have low baseline firing rates but are robustly activated by levodopa. In contrast, D2-MSNs may be undersampled because their low baseline firing rates limit detection of further suppression. Second, while fiber photometry enables longitudinal monitoring of the same ensemble of cells, electrophysiological recordings often sample different neurons and mice across timepoints, given that single units can be lost or gained over days in a given animal, and recording quality declines over time after implantation. These factors may increase the variability of the electrophysiology data. Despite these limitations in detecting activity suppression, our findings suggest that stronger D2-MSN suppression may also reflect an increase in the number of suppressed cells or alterations in non-somatic calcium dynamics^77^, rather than firing rate alone.

As with the changes in D1-MSNs, stronger suppression of D2-MSNs observed by day 4 could be mediated by dopamine signaling, intrinsic or synaptic properties. However, we did not find alterations in D2 receptor sensitivity, intrinsic excitability, or excitatory synaptic transmission during this early phase of treatment. An alternative mechanism would be increased synaptic inhibition of D2-MSNs. Several sources provide inhibitory inputs to D2-MSNs^78–80^, however, given that D2-MSN suppression parallels heightened D1-MSN activity, direct lateral inhibition from D1-MSNs emerges as a strong candidate. Increased levodopa-evoked D1-MSN firing over the course of treatment could drive the increased inhibition of D2-MSNs even without changes in the strength of these synapses. In turn, decreased D2-MSN activity would further disinhibit D1-MSNs. Consistent with this possibility, we previously demonstrated that in parkinsonian mice, selective D1R agonists can decrease putative D2-MSNs firing^10^, and we have recently shown that lateral MSN-MSN connections are key drivers of LID^11^.

In addition to the more rapid and severe onset of LID, we observed that the LID offset hastened between day 4 and week 4 of treatment. Interestingly, this is not linked to changes in D1-MSN activity, but rather to a faster recovery of D2-MSN activity. The concomitant reduction in D2 receptor sensitivity may contribute to this faster rebound, although other mechanisms, including changes in D2 receptor internalization^81^ and state-dependent changes in glutamatergic transmission^22^, may also contribute. Importantly, these results suggest that the more rapid LID offset is not a passive consequence of declining dopamine levels, but reflects active circuit-level processes, in this case a faster rebound in D2-MSN activity. These observations may have broader implications for understanding other motor complications that have traditionally been attributed primarily to striatal dopamine levels and buffering capacity^82^.

Some changes in MSNs properties may reflect homeostatic adaptations, such as the reduction in D1-MSN intrinsic excitability, consistent with a previous report^14^. This decrease in excitability developed between day 4 and week 4, after the increase in excitatory inputs had already occurred and without corresponding change in D1-MSN responses, potentially reflecting a compensatory response. This change was accompanied by a larger AHP and a reduction in spike half-width. While we did not directly measure ionic conductances, these changes are consistent with an increase in BK and/or SK currents. Additionally, our analysis using subthreshold current injections suggests upregulation of A-type potassium currents. Because BK, SK, and I_A_ are regulated by endogenous dopamine via D1 receptors^34^, their increased availability may facilitate dopamine modulation during the “ON” state. Thus, while upregulation of potassium currents may be a homeostatic reaction to sustained elevations in dopamine signaling, it may be maladaptive. Indeed, despite reduced D1-MSN intrinsic excitability after 4 weeks, peak levodopa-evoked firing rates and GCaMP signals were similar to day 4.

Prevailing models of basal ganglia function propose that co-activation of D1- and D2-MSNs facilitates desired actions while suppressing competing ones^25^. This model predicts that achieving antiparkinsonian effects without dyskinesia requires restoring D1- and D2-MSN co-activation. Our analysis of D1- and D2-MSN activity across behavioral states support this prediction: during prokinetic periods without dyskinesia, both D1-MSN and D2-MSN activity remained elevated, while during LID periods, D1-MSN activity remained elevated but D2-MSN activity was markedly suppressed (Figure S4A). This suppression may be crucial for the transition from antiparkinsonian effects to LID, by disrupting co-activation, and thus impairing the ability to inhibit competing movements. Overall, these findings reinforce the idea that both striatal populations play essential roles in LID pathophysiology and that excitatory and inhibitory synaptic connections are likely to contribute to the evolution of LID over the course of levodopa treatment in PD.

## RESOURCE AVAILABILITY

### Lead contact

Requests for further information and resources should be directed to the Lead contact, Alexandra Nelson.

### Materials availability

This study did not generate any new unique materials.

### Data and code availability

All data and code generated from this publication are available on Zenodo (DOI: 10.5281/zenodo.23677819). Any additional information required to reanalyze the data reported in this paper is available from the lead contact upon request. An earlier version of this manuscript was posted to bioRxiv on May 21, 2025 at doi.org/10.1101/2025.05.16.654598.

## ACKNOWLEDGEMENTS

This work was supported by grants from the Parkinson’s Foundation (PF-PRF-1045114 to R.M.P.), the National Institute of Neurological Disorders and Stroke (NINDS) (R01NS101354 to A.B.N. & K99NS149250 to R.M.P), Richard and Shirley Cahill Endowed Chair in Parkinson’s Disease Research (to A.B.N.). Additionally, this research was funded in part by Aligning Science Across Parkinson’s (ASAP020529 to A.B.N.) through the Michael J. Fox Foundation for Parkinson’s Research (MJFF). For the purpose of open access, the author has applied a CC BY public copyright license to all Author Accepted Manuscripts arising from this submission. We thank all the members of the Nelson Lab for helpful feedback.

## AUTHOR CONTRIBUTIONS

Conceptualization, R.M.P., M.B.R. & A.B.N.; Methodology, R.M.P., M.B.R. & A.B.N.; Investigation, all authors; Formal Analysis, R.M.P., P.M., V.D. & A.B.N.; Writing – Original Draft, R.M.P. & A.B.N.; Writing – Review & Editing, all authors; Visualization: R.M.P. & A.B.N.; Funding Acquisition, R.M.P. & A.B.N.; Supervision, A.B.N.

## DECLARATION OF GENERATIVE AI AND AI-ASSISTED TECHNOLOGIES IN THE MANUSCRIPT PREPARATION PROCESS

During the preparation of this work, the authors used ChatGPT for MATLAB analysis code generation and error identification. The authors reviewed the content as needed and take full responsibility for the content of the publication.

## STAR METHODS

### EXPERIMENTAL MODEL AND STUDY PARTICIPANT DETAILS

#### Animals

All animal procedures were approved by the University of California San Francisco Institutional Animal Care and Use Committee (IACUC). Animals were housed under a 12h light/dark cycle with access to food and water ad libitum. For all experiments, both male and female C57Bl/6J mice, aged 3-12 months, were used. For *ex vivo* electrophysiology experiments and c-Fos staining, D1-tdTomato or D2-GFP mice were used. The following mouse lines were used for *in vivo* experiments: GCaMP fiber photometry: D1-Cre and A2a-Cre; GRAB-DA fiber photometry: wild type; *in vivo* single-unit electrophysiology: D1-Cre, A2a-Cre and wild type. Age-matched littermates were randomly assigned to experimental and control groups.

### METHOD DETAILS

#### Surgical Procedures

A detailed surgical protocol can be found at (DOI: dx.doi.org/10.17504/protocols.io.dm6gpdw1dgzp/v1). Briefly, three- to six-month old mice were anesthetized with a combination of ketamine/xylazine (40/10 mg/kg IP) and inhaled isoflurane (1%). After placement in the stereotaxic frame (Kopf Instruments), the scalp was opened, and a mounted drill was used to create a hole over the left medial forebrain bundle (MFB). Using a 33-gauge needle (WPI) or a syringe with removable glass needle (Hamilton) and Micro4 pump (WPI), 1 μL of 6-hydroxydopamine (6-OHDA, Sigma-Aldrich, 5 μg/μl in normal saline) was injected unilaterally into the MFB (-1.0 AP, -1.0 ML, -4.9 DV) at a rate of 0.2 μl/min. To reduce toxicity to other monoaminergic axons, desipramine (Sigma-Aldrich, 25mg/kg IP) was administered 30 min prior to 6-OHDA injection. After the surgery, animals received daily saline injections and were provided high-fat dietary supplements (Diet-Gel, peanut butter, forage mix) for a 1-2 week recovery period.

Viral injections were performed during the same surgical procedure as the MFB injection described above. A mounted drill was used to create a hole over the left dorsolateral striatum (DLS, +0.8 AP, - 2.4 ML, -3 DV). For *ex vivo* electrophysiology experiments with GIRK overexpression, D2-GFP mice were injected with 500 nL AAV9-hsyn-TdTomato-T2A-mGIRK2-1-A22A-WPRE-bGH (titer: 2.7e13/ml, undiluted, UPenn Vector Core) into the DLS. For fiber photometry experiments, D1-Cre or A2A-Cre mice were injected with 500 nL of a Cre-dependent GCaMP6s (original titer: 3.056e13/ml, diluted 1:9, UPenn Vector Core) into the DLS, whereas wild type mice were injected with 500 nL of AAV9-hsyn-GRAB-DA2h (titer: 1.88e13/ml, undiluted, Addgene 140554) and implanted with a fiber-optic ferrule (0.4 mm, Doric Lenses) above the DLS (-2.7 DV). Fibers were secured with dental cement (Metabond) and dental acrylic (Ortho-Jet or Henry Schein).

For *in vivo* single-unit electrophysiology experiments, the scalp was reopened 3-4 weeks after the initial surgeries to implant the recording device. Two different recording devices were used: either a fixed 32-channel electrode array (Innovative Neurophysiology) or a high density 64-channel silicon probe (Cambridge Neurotech). For 32-channel electrode array implantation, three additional holes were drilled for two skull screws (FST) and a ground wire. The array was slowly inserted through the craniectomy into the DLS (-2.5 DV). A thin layer of dental cement (Metabond) was applied to the surface of the skull and dental acrylic (Ortho-Jet or Henry Schein) was applied to cover all exposed hardware. For silicon probe implantation, a 2-shank 64-channel high-density silicon probe (model H6, Cambridge Neurotech), was mounted on a microdrive and implanted above the DLS, and a ground wire above the cerebellum. A head bar was also attached to the skull to allow tethering without damaging the probe. The recording electrodes were stained with DiI or DiO lipophilic dyes (Thermo Fisher) prior to implantation for post hoc identification of the electrode track. The experiments were performed at least one day after implantation. For some timepoints (like day 3-5 and week 4), where multiple days were recorded, the probe depth was finely adjusted by moving the microdrive so that different neurons were recorded each day.

#### Behavior

Details of the behavioral assessments can be found at (DOI: https://www.protocols.io/view/behavioral-testing-open-field-and-dyskinesia-scori-6qpvr67oovmk/v1). Briefly, after a 6-week post-surgical baseline period, animals began daily treatment. Animals were placed on an open field arena (25 cm diameter cylinder) and video recorded with a side and a top camera. Gross movement was quantified using video-tracking software (Noldus Ethovision). Parkinsonian animals receiving levodopa developed robust levodopa-induced dyskinesia (LID). Dyskinesia was quantified by the experimenter using the Abnormal Involuntary Movement score (AIMs), a standardized metric that considers dyskinesia across axial, limb, and orolingual body segments. As described by Cenci and Lundblad (2007), dyskinesia severity in each body segment was quantified during 60 s observation windows as follows: 0 = normal movement, 1 = abnormal movement for < 50% of the time, 2 = abnormal movement for > 50% of the time, 3 = abnormal movement for 100% of the time, but can be interrupted, and 4 = continuous, uninterruptable abnormal movement. The total AIM score represents the summation of the AIM score for each body segment (axial, limb, orolingual), for a maximum possible score of 12. For *ex vivo* experiment animals, AIMs were scored every 20 minutes for a 2-hour period. For *in vivo* experiments (GCaMP & GRAB-DA fiber photometry and single-unit electrophysiology), AIMs were scored every minute for the first 20 minutes after injection, and every 5 minutes thereafter to complete 120 minutes.

#### Pharmacology

Details for the preparation of pharmacological agents can be found at (DOI: dx.doi.org/10.17504/protocols.io.e6nvw1nbzlmk/v2). For all the experiments, 5 mg/kg levodopa (Sigma-Aldrich) was co-administered with 2.5 mg/kg benserazide (Sigma-Aldrich) and prepared in normal saline. Levodopa was given via IP injection 5 days per week. For *in vivo* experiments, daily levodopa injections continued for 4 weeks. For *ex vivo* experiments, levodopa was administered either for 3 days or for 4 weeks before the experiment. Picrotoxin (Sigma-Aldrich) was dissolved in warm water at 5 mM and added to ACSF for a final concentration of 50 μM. SCH23390 (Tocris) was dissolved in water at 10 mM and added to ACSF for a final concentration of 10 μM. AP5 (Abcam) was dissolved in water at 50 mM and added to ACSF for final concentrations of 50 μM. Scopolamine (Tocris) was dissolved in water at 200 μM and added to ACSF for final concentrations of 200 nM.

CGP55845 (Tocris) was dissolved in water at 300 μM and added to ACSF for final concentrations of 300 nM. Dopamine (Sigma-Aldrich) was dissolved in water with 0.1% ascorbic acid at 1 or 100 mM and added to ACSF for a final concentration of 1 or 100 μM. CNQX (Sigma-Aldrich or Tocris) was dissolved in water at 10 mM and added to ACSF for a final concentration of 10 μM. TTX (Tocris) was dissolved in water at 1 mM and added to ACSF for a final concentration of 1 μM. Sulpiride (Tocris) was dissolved in water at 10 mM and added to ACSF for a final concentration of 10 μM.

#### Fiber photometry

A detailed protocol for fiber photometry can be found at https://doi.org/10.17504/protocols.io.8epv59dbjg1b/v1. Animals were habituated to IP injections and tethering for at least 30 minutes/ session for 2 sessions before starting the experiments. Mice were placed in a circular open field (25 cm diameter) and recorded for a baseline period of 30 minutes before saline or levodopa injection. For saline sessions, GCaMP signals were recorded for 1 hour after injection. For levodopa sessions, GCaMP signals were recorded for 2 hours after injection. A side camera and a top camera (20 fps, Imaging Source) allowed for post-hoc quantification of gross movement using an offline video-tracking software (Noldus Ethovision). Fine behavior was manually scored by the experimenter (AIM score). Fiber photometry signals were acquired through implanted 400 μm optical fibers, using an LED driver system (Doric). The LED excitation level was determined empirically in a pilot study by selecting the lowest power that provided an adequate signal-to-noise ratio while minimizing photobleaching. The LED power was set the same across all recordings. In our fiber photometry system (TDT), the LEDs are frequency modulated at 210 Hz (for 405 nm) and 330 Hz (for 465 nm). The resulting power at the fiber tip (measured with sinusoidal excitation) was: 3.3 µW for the 405 nm LED and 10 µW for the 465 nm LED. Following signal modulation, 405 nm (isosbestic) and 465 nm signals were demodulated via a lock-in amplifier (RZ5P, TDT), visualized, and recorded (Synapse, TDT). Videos were synchronized with simultaneous photometry recordings via TTL pulses triggered by the video tracking software. IP injection times were synchronized with simultaneous photometry recordings via manually triggered TTL pulses.

#### *In vivo* electrophysiology

*In vivo* extracellular recordings were performed using two different recording devices: a fixed 32-channel microwire (35 mm tungsten) array (Innovative Neurophysiology) or a drivable high-density silicon probe (Cambridge Neurotech chronic 64 channel H6, 2 shanks, 9 mm length). A detailed protocol for *in vivo* electrophysiology can be found at: dx.doi.org/10.17504/protocols.io.36wgq641ylk5/v1 (for Cambridge Neurotech silicon probes) and at: dx.doi.org/10.17504/protocols.io.b9ucr6sw (Innovative Neurophysiology electrode arrays). Briefly, mice were habituated to the open field (25 cm diameter), tethering and IP injections (saline). The animal’s gross behavior was recorded by video tracking software (Noldus Ethovision). Fine behavior was manually scored by the experimenter (AIM score). Recording sessions consisted of a 30-minute baseline period, followed by IP injection of levodopa. Electrophysiological recordings continued for at least 80 minutes post-injection. Behavioral measurements were synchronized with simultaneous electrophysiological recordings via TTL pulses triggered by the video tracking software and recorded by the electrophysiology system.

For recordings using fixed a 32-channel microwire array the animal was placed in the open-field while tethered via a lightweight, multiplexed headstage cable (Triangle Biosystems) attached to a low-torque electrical commutator (Dragonfly) to allow free movement. Single unit activity from microwires was recorded using a 32-channel recording system (MAP system, Plexon). Spike waveforms were filtered at 154–8800 Hz and digitized at 40 kHz. The experimenter manually set a threshold for storage of electrical events. To verify the location of the optrode array in DLS, mice were deeply anestethized after the last recording session and, prior to transcardia perfusion, electrolytic lesions were made to mark electrode tips using a solid state, direct current Lesion Maker (Ugo Basile), by applying 100 μA for 5 s per microwire. Mice were then terminally anesthetized with ketamine/xylazine (200/40 mg/kg i.p.), transcardially perfused with 4% paraformaldyde (PFA), and the brain dissected from the skull. The brain was post-fixed overnight in 4% PFA and then placed in 30% sucrose at 4°C for histology processing. A subset of these recordings were reported previously, with a different scientific question and analysis approach^10,13^.

For high-density silicon probe recordings, we recorded the signals at 30 kHz using an INTAN system (RHD2000 USB Interface Board, INTAN Technologies). Animals were tethered via an ultra-thin SPI interface cable (INTAN Technologies) attached to an electrical commutator (Doric). Signals were digitized at 30 kHz. To verify the location of the probe tracks, mice were terminally anesthetized with ketamine/xylazine, perfused with 4% PFA, and the brain dissected for histology processing.

#### *Ex vivo* electrophysiology

A detailed protocol for *ex vivo* electrophysiology can be found at (DOI: dx.doi.org/10.17504/protocols.io.kqdg3k6n1v25/v1). To prepare acute brain slices, mice were anesthetized with ketamine/xylazine and transcardially perfused with 95%O_2_/5%CO_2_ oxygenated, ice-cold glycerol-based artificial cerebrospinal fluid (ACSF) containing (in mM): 250 glycerol, 2.5 KCl, 1.2 NaH_2_PO_4_, 10 HEPES, 21 NaHCO_3_, 5 D-Glucose, 2 MgCl_2_, 2 CaCl_2_. For experiments designed to simulate a physiological “ON” state, animals received IP injections of levodopa-benserazide (5mg/kg-2.5mg/kg) 30 minutes before decapitation. Following decapitation, brains were dissected and sequential coronal slices (275 μm) containing the striatum were collected with a vibratome (Leica). Slices were immediately transferred to a holding chamber containing warm (33°–34°C), carbogenated ACSF containing (in mM): 125 NaCl, 26 NaHCO_3_, 2.5 KCl, 1.25 NaH_2_PO_4_, 12.5 D-Glucose, 1 MgCl_2_, 2 CaCl_2_. Slices were incubated for 45 min and then held at room temperature (22°–24°C) until used for recording.

For all recordings, slices were transferred to a recording chamber superfused (∼2 mL/min) with carbogenated ACSF (31°–33°C). Medium spiny neurons were targeted for recordings using differential interference contrast (DIC) optics on an Olympus BX 51 WIF microscope. In Drd1a-tdTomato mice, D1-MSNs were identified by their tdTomato-positive somata. Conversely, D2-MSNs were identified by their medium-sized tdTomato-negative somata. In D2-GFP mice, D2-MSNs were identified by their GFP-positive somata and D1-MSNs were identified by their GFP-negative, medium-sized somata. Fluorescent-negative neurons with interneuron-like physiological properties were excluded from the dataset.

Medium spiny neurons were patched in the whole-cell configuration using borosilicate glass electrodes (2-5 MΩ). For voltage-clamp measurements of miniature excitatory postsynaptic currents (mEPSC) at a holding potential of -70mV, electrodes were filled using a cesium-based internal solution containing (in mM): 120 CsMeSO3, 15 CsCl, 8 NaCl, 0.5 EGTA, 10 HEPES, 2 MgATP, 0.3 NaGTP, 5 QX-314, pH 7.3. Picrotoxin (50 μM) and TTX (1 μM) was added to the external solution. For current-clamp experiments, electrodes were filled using a K-based internal solution containing (in mM): 130 KMeSO_3_, 10 NaCl, 2 MgCl_2_, 0.16 CaCl_2_, 0.5 EGTA, 10 HEPES, 2 MgATP, 0.3 NaGTP, pH 7.3). Picrotoxin (50 μM) was added to the external solution. After whole-cell break in, cells were given at least 5 min to dialyze internal solution before recording. Whole-cell recordings were made using a MultiClamp 700B amplifier (Molecular Devices) and digitized with an ITC-18 A/D board (HEKA). Data were acquired using Igor Pro 6.0 software (Wavemetrics) and custom acquisition routines (mafPC, courtesy of M.A. Xu-Friedman). For voltage-clamp measurements of dopamine-evoked GIRK currents, whole-cell recordings were made from doubly fluorescent (GFP^+^ and tdTomato^+^;GIRK2^+^) MSNs. Electrodes were filled using an internal solution containing (in mM): 135 mM D-Gluconic Acid (K), 2 mM MgCl_2_, 0.1 mM CaCl_2_, 10 mM HEPES (K), 10 mM BAPTA-tetrapotassium, 1 mg/ml ATP, 0.1 mg/ml GTP, and 1.5 mg/ml phosphocreatine (pH 7.3, 285 mOsm). GFP^+^/GIRK2^+^ MSNs were held at −60 mV. Picrotoxin (50 μM), SCH23390 (10 μM), AP5 (50 μM), Scopolamine (200 nM), CGP55845 (300 nM) and CNQX (10 μM) were always present in the external solution to isolate D2R-mediated increases in GIRK currents. Dopamine was applied at 1 μM by bath perfusion. After ∼3-4 minutes dopamine was washed out with ACSF for 4-5 minutes and then 100 μM dopamine was applied. For experiments designed to simulate a physiological “ON” state, recordings were restricted to the 4-hour period after levodopa injection.

#### Histology and microscopy

A detailed protocol for preparation of histological sections can be found at (DOI: dx.doi.org/10.17504/protocols.io.b9ubr6sn). After behavioral experiments, mice were anesthetized with ketamine/xylazine and transcardially perfused with 4% paraformaldehyde (PFA) in phosphate buffer solution (PBS). Following perfusion, brains were dissected and post-fixed overnight in 4% PFA, then transferred to 30% sucrose and stored at 4°C for cryoprotection. Brains were then sliced coronally into 30 μm sections on a freezing microtome (Leica).

After *ex vivo* electrophysiology experiments, 275 μm slices were stored overnight in 4% PFA. The following day, slices were transferred to 30% sucrose and sub-sectioned into 50 μm sections on a freezing microtome (Leica).

For immunohistochemistry, the tissue was blocked in 3% normal donkey serum (NDS) and permeabilized with 0.1% Triton X-100 on a room-temperature (RT) shaker. Primary anti-bodies were used in the appropriate combination: Rabbit anti-TH (Pel-Freez Biologicals, 1:1000) or chicken anti-TH (Milipore, 1:1000); mouse anti-RFP (Rockland, 1:1000); rabbit anti c-Fos (Cell Signaling Technology, 1:1000); goat anti-ΔfosB (Santa Cruz Biotechnology, 1:1000) were added to the NDS and tissue was incubated overnight on a 4°C shaker. Tissue was then incubated in appropriate secondary antibodies from Jackson ImmunoResearch Labs (Donkey anti-rabbit Alexa Fluor 488, 594 or 647, 1:500; Donkey anti-mouse Alexa Fluor 594, 1:500; Donkey anti-goat Alexa Fluor 488, 1:500; Donkey anti-chicken Alexa Fluor 647, 1:500) either overnight (16 h) on a 4°C shaker or for 2 hours at RT before being washed and mounted (Vectashield Mounting Medium) onto glass slides for imaging. Images (4x or 10x) were obtained using a Nikon 6D conventional widefield microscope or an Olympus Fluoview FV3000 confocal microscope. In 6-OHDA-treated mice, the extent of dopamine depletion was confirmed by TH immunohistochemistry. Adequate viral expression (GCaMP, GRAB-DA, GIRK-tdTomato) was confirmed.

### QUANTIFICATION AND STATISTICAL ANALYSIS

#### Statistics

Details can be found in Table S1. Statistical tests were performed using GraphPad Prism 10. All data are presented as the mean ± SEM (unless specified otherwise in the figure legends) with “n” referring to the number of cells and “N” referring to the number of animals.

#### Behavior

Dyskinesia was quantified using the Abnormal Involuntary Movement score (AIMs), as described above. The motion of the mouse was tracked using an offline video tracker software (Noldus, Ethovision), that tracked the center of gravity of the animal in the open field arena. Mouse velocity was quantified as the average velocity in 1-minute bins (Figure S1). Changes in AIMs and velocity (Figure 1 and S1), were compared across treatment timepoints using repeated-measures one-way ANOVA, with a post-hoc Tukey test.

#### Fiber photometry

Offline, the 405 nm signal was fit to the 465 nm signal using a first-degree polynomial fit (using Matlab) to extract the non-calcium-dependent signal (due to autofluorescence, fiber bending, etc.). The fitted 405 nm signal was then subtracted from the 465 nm signal to generate a motion-corrected signal. For GCaMP experiments, we corrected for the gradual bleaching observed over the ∼2.5-hour recordings by fitting an exponential curve to the signal and normalizing it by the fit. The resulting signal was then smoothed to reduce noise, by taking the moving average over a window of 0.5s. To detect GCaMP transients, we calculated the envelope of the ΔF/F signal to capture meaningful fluctuations, and identified peaks and valleys of the envelope. Transients were defined as valley-to-peak difference exceeding 2% ΔF/F. The baseline-subtracted transient count or ΔF/F values were binned in 1-minute intervals and averaged across animals (Figure 2; Figure S3). Changes in ΔF/F or ΔTransients (Figures 2; Figure S2) across treatment timepoints were compared using repeated-measures one-way ANOVA, with a post-hoc Tukey test. The linear correlation between ΔF/F or ΔTransients and AIMs was calculated using Pearson correlation. For GRAB-DA experiments, ΔF/F was extracted, binned into 1-minute intervals, and aligned to levodopa injection (Figure 5). Changes in peak ΔF/F across treatment timepoints were compared using repeated-measures one-way ANOVA, with a post-hoc Tukey test.

To quantify the D1-MSN and D2-MSN GCaMP transient frequency associated with behavioral states (Figure S4), we calculated the average transient frequency for each mouse across all recording sessions during three behavioral states: pre-levodopa, post-levodopa prokinetic phase (defined as periods following levodopa administration when locomotor velocity exceeded 1.5 cm/s and AIMs = 0), and post-levodopa dyskinesia (defined as periods with AIMs > 1). Transient frequency during the dyskinesia phase was then stratified by AIM severity.

#### *In vivo* electrophysiology

For recordings performed using 32-channel microwire arrays (Innovative Neurophysiology) and an associated 32-channel recording system (MAP system, Plexon), single units (SUs) were identified offline by manual sorting into clusters (Offline Sorter, Plexon). Waveform features used for separating units were typically a combination of valley amplitude, the first three principal components (PCs), and/or nonlinear energy. Clusters were classified as SUs if they fulfilled the following criteria: (1) < 1% of spikes occurred within the refractory period and (2) the cluster was statistically different (p < 0.05, MANOVA using the aforementioned features) from the multi- and other single-unit clusters on the same wire. For recordings performed with 64-channel high-density probes and INTAN system, we classified SUs by automated spike sorting (KiloSort, https://github.com/cortex-lab/Kilosort). SUs were then manually curated using Phy (https://github.com/cortex-lab/phy). SUs were then classified as putative medium spiny neurons (MSNs) as previously described^10,55–57^, using features of the spike waveform (peak to valley and peak width), as well as inter-spike interval distribution. Only putative MSNs were included in subsequent analyses.

Firing rate was averaged in 1-minute bins. Modulation of firing rate by levodopa was determined by comparing SU firing rates before and after levodopa administration. The 30-minute baseline period was compared to a 30-minute period following drug injection (10-40 minutes post-injection). Following levodopa administration, SUs were categorized into three broad groups as follows, based on significant changes in firing rate (p < 0.01, Wilcoxon rank-sum test (denoted Mann-Whitney)) following levodopa treatment: putative D1-MSNs (increase in firing rate), putative D2-MSNs (decrease in firing rate), or no change units (nonsignificant change in firing rate). Putative D1-MSNs with a significant correlation (and Pearson correlation coefficient r>0.4) were classified as LID-associated putative D1-MSNs. Putative D2-MSNs with a significant correlation (and Pearson correlation coefficient r<0) were classified as LID-associated putative D2-MSNs.

Changes in the levodopa-evoked increases or decreases in firing rates across treatment timepoints were compared using a Kruskal-Wallis test with a Dunn’s post-hoc (Figure 4; Figure S5).

#### *Ex vivo* electrophysiology

To characterize intrinsic excitability, custom code in Igor Pro (Wavemetrics) was used to extract the instantaneous firing rate for each current step, rheobase, maximum firing rate, resting membrane potential, action potential threshold, spike half width, afterhyperpolatization (AHP) and voltage deflections to depolarizing and hyperpolarizing current steps. The number of action potentials for every step was manually counted. For excitability, current-response curves were compared using a two-way repeated measures ANOVA across treatment timepoints and a Geissner-Greenhouse correction, with a post-hoc Tukey test. For comparing passive and active properties across treatment timepoints, significance was determined using a Kruskal-Wallis test. A Dunn’s post hoc test was applied to test for significance between individual groups (Figure 7; Figure S8 & S9). For mEPSC frequency and amplitude measurements, custom code in Igor Pro (Wavemetrics) was used to extract mEPSCs events. Only cells with at least 500 detected events were included in subsequent analysis. Cumulative probability plots were generated from the first 500 mEPSC events per cell. The average amplitudes and frequencies for each cell were compared across treatment timepoints using a Kruskal-Wallis test with a Dunn’s post-hoc (Figure 6 & S7). Dopamine-evoked GIRK currents were quantified manually using Igor Pro and Matlab. A Kruskal-Wallis test with a Dunn’s post-hoc was used to compare the amplitudes and ratio of dopamine-evoked GIRK currents (Figure S10).

#### Immunohistochemistry quantification

c-Fos^+^ cells were detected from coronal striatal sections using QuPath (open-source software for image analysis, https://qupath.github.io/). Detailed methods for quantification can be found at: https://github.com/UCSF-Nelson-Lab/QuPath-scripts-for-cFos-analysis (doi.org/10.5281/zenodo.21685413). Briefly, images were imported into QuPath. The DLS region was manually annotated for 2-3 sections per mouse. c-Fos^+^ or ΔFosB^+^ cell bodies were identified and quantified using a custom script, which was adapted from the extension StarDist from QuPath. c-Fos immunoreactivity in the dopamine-intact striatum was minimal; however, ΔFosB immunoreactivity was detectable. To account for potential non-specific ΔFosB labeling, cells in the dopamine-depleted striatum were considered positive only if their fluorescence intensity exceeded the 95% confidence interval of the intensity distribution measured in the dopamine-intact striatum. Colocalization of c-Fos^+^/ΔFosB^+^ cell bodies with tdTomato was performed with a custom script adapted from https://gist.github.com/Svidro/68dd668af64ad91b2f76022015dd8a45. The custom-made scripts can be found at: https://github.com/UCSF-Nelson-Lab/QuPath-scripts-for-cFos-analysis. Changes in the number of c-Fos^+^/ΔFosB^+^ and double c-Fos^+^/tdTomato^+^ or ΔFosB^+^/tdTomato^+^ cells across treatment timepoints were compared using a Kruskal-Wallis test with a Dunn’s post-hoc (Figure 3). The degree of striatal dopamine loss was quantified by measuring the percentage of remaining TH staining relative to the dopamine-intact contralateral side (Figure S11). Specifically, the background-subtracted full-field fluorescence TH signal in the dorsolateral striatum ipsilateral to the MFB lesion was normalized to the corresponding signal in the contralateral striatum. The average TH signal measured in the medial corpus callosum was used as background reference.

### KEY RESOURCES TABLE

| REAGENT or RESOURCE | SOURCE | IDENTIFIER |
| --- | --- | --- |
| **Antibodies** |  |  |
| Rabbit anti-TH | Pel-Freez | RRID: AB_2617184; Cat# P40101-150 |
| Chicken anti-TH | Milipore | RRID: AB_570923; Cat# AB9702 |
| Mouse anti-RFP | Rockland | RRID: AB_2611063; Cat# 200-301-379 |
| Rabbit anti c-Fos | Cell Signaling Technology | RRID: AB_2247211; Cat# 2250 |
| Goat anti-deltaFosB | Santa Cruz Biotechnology | RRID: AB_631516; Cat# sc-48-G |
| Alexa Fluor 488 Donkey Anti-Rabbit IgG | Jackson ImmunoResearch | RRID: AB_2340619; Cat# 711-546-152 |
| Alexa Fluor 594 Donkey Anti-Rabbit IgG | Jackson ImmunoResearch | RRID: AB_2340621; Cat# 711-585-152 |
| Alexa Fluor 647 Donkey Anti-Rabbit IgG | Jackson ImmunoResearch | RRID: AB_2534017; Cat# A10042 |
| Alexa Fluor 488 Donkey Anti-Goat IgG | Jackson ImmunoResearch | RRID: AB_2336933; Cat# 705-545-147 |
| Alexa Fluor 594 Donkey Anti-Mouse IgG | Jackson ImmunoResearch | RRID: AB_2340857; Cat# 715-586-150 |
| Alexa Fluor 647 Donkey Anti-Chicken IgY (IgG) | Jackson ImmunoResearch | RRID: AB_2340380; Cat# 703-606-155 |
| **Bacterial and virus strains** |  |  |
| AAV9-hsyn-TdTomato-T2A-mGIRK2-1-A22A-WPRE-bGH | UPenn | V4822MI-R |
| AAV9-hsyn-GRAB-DA2h | Yulong Li Lab | RRID: Addgene_140554; lot v188117 |
| AAV1.Syn.Flex.GCaMP6s.WPRE.SV40 | UPenn | RRID: Addgene_100845; lot CS1323 |
| **Chemicals, peptides, and recombinant proteins** |  |  |
| DiO | Invitrogen | Cat# V22889 |
| 6-Hydroxydopamine hydrobromide | Sigma-Aldrich | Cat# H116 |
| Despiramine hydrochloride | Sigma-Aldrich | Cat# D3900 |
| Benserazide hydrochloride | Sigma-Aldrich | Cat# B7283 |
| 3,4-Dihydroxy-L-phenylalanine | Sigma-Aldrich | Cat# D9628 |
| Lidocaine N-ethyl chloride | Sigma-Aldrich | Cat# L1663 |
| Cesium methanesulfonate | Sigma-Aldrich | Cat# C1426 |
| Potassium Methanesulfonate | Sigma-Aldrich | Cat# 83000 |
| Potassium D-Gluconate | Sigma-Aldrich | Cat# G4500 |
| BAPTA-Na<sub>4</sub> | Sigma-Aldrich | Cat# 14513 |
| Phosphocreatine disodium salt hydrate | Sigma-Aldrich | Cat# P7936 |
| Guanosine 5'-triphosphate sodium salt | Sigma-Aldrich | Cat# G8877 |
| Adenosine 5'-triphosphate magnesium salt | Sigma-Aldrich | Cat# A9187 |
| Picrotoxin | Sigma-Aldrich | Cat# P1675 |
| CNQX | Tocris | Cat# 1045 |
| CNQX | Sigma-Aldrich | Cat# C239 |
| DL-APV | Abcam | Cat# AB120271 |
| Triton X-100 | Sigma-Aldrich | Cat# T8787 |
| Tetrodotoxin | Tocris | Cat# 1069 |
| SCH23390 | Tocris | Cat# 0925 |
| Scopolamine | Tocris | Cat# 1414 |
| CGP55485 | Tocris | Cat# 1248 |
| Dopamine | Sigma-Aldrich | Cat# H8502 |
| **Critical commercial assays** |  |  |
| VECTASHIELD Antifade Mounting Medium | Vector Laboratories | RRID: AB_2336789; Cat# H-1000 |
| **Experimental models: Organisms/strains** |  |  |
| Mouse: Wild-type C57Bl/6J | The Jackson Laboratory | RRID: IMSR_JAX:000664 |
| Mouse: Stock Tg(Drd2-EGFP)S118Gsat/Mmnc Mus Musculus | MMRRC | RRID: MMRRC_000230-UNC |
| Mouse: B6.FVB(Cg)-Tg(Drd1-Cre)EY217Gsat/Mmucd | MMRRC | RRID: MMRRC_034258-UCD |
| Mouse: B6.FVB(Cg)-Tg(Adora2a-Cre)KG139Gsat/Mmucd | MMRRC | RRID: MMRRC_036158-UCD |
| Mouse: FVB.Cg-Tg(Drd1a-tdTomato)5Calak/Mmnc | MMRRC | RRID: MMRRC_030512-UNC |
| **Deposited data** |  |  |
| Dataset and code | Zenodo | 10.5281/zenodo.23677819 |
| Code to count cFos/fosB | Zenodo | 10.5281/zenodo.21685413;<br>https://github.com/UCSF-Nelson-Lab/QuPath-scripts-for-cFos-analysis |
| Code for GCaMP data extraction | Zenodo | 10.5281/zenodo.21686313;<br>https://github.com/UCSF-Nelson-Lab/photometry |
| Code for GRAB-DA Analysis | Zenodo | 10.5281/zenodo.21687408;<br>https://github.com/UCSF-Nelson-Lab/GRAB-DA_Analysis |
| Code for in vivo electrophysiology analysis | Zenodo | 10.5281/zenodo.21687765;<br>https://github.com/UCSF-Nelson-Lab/InVivo-SingleUnit-Analysis |
| In vivo electrophysiology data | DANDI Archive | RRID: SCR_017571;<br>DOI:<br>https://doi.org/10.48324/dandi.001180/0.261024.0010 |
| **Software and algorithms** |  |  |
| IgorPro | Wavemetrics | RRID: SCR_00325;<br>http://www.wavemetrics.com/products/igorpro/igorpro.htm |
| mafPC (software package for use with IgorPro) | Xu-Friedman Lab | https://www.xufriedman.org/mafpc |
| MATLAB R2015a | MathWorks | RRID: SCR_001622;<br>https://www.mathworks.com/products/matlab/ |
| Adobe Illustrator | Adobe | RRID: SCR_014198;<br>http://www.adobe.com/products/illustrator |
| ImageJ | NIH | RRID: SCR_003070;<br>https://imagej.nih.gov/ij/ |
| EthovisionXT | Noldus | RRID: SCR_000441;<br>http://www.noldus.com/animal-behavior-research/products/ethovision-xt |
| MAP Software (RASPUTIN 2.4) | Plexon | RRID: SCR_003170;<br>https://plexon.com/wp-content/uploads/2017/06/RASPUTIN-Manual.pdf |
| Offline Sorter | Plexon | RRID: SCR_000012;<br>http://plexon.com/products/offline-sorter |
| NeuroExplorer | Plexon | RRID: SCR_001818;<br>http://www.neuroexplorer.com/ |
| Axon MultiClamp Commander | Axon | RRID: SCR_018455;<br>http://mdc.custhelp.com/app/answers/detail/a_id/18877/~/axon%E2%84%A2-multi |
| NIS-Elements | Nikon | RRID: SCR_014329;<br>https://www.microscope.healthcare.nikon.com/products/software |
| NeuroInfo | MBF Biosciences | RRID: SCR_017346;<br>https://www.mbfbioscience.com/products/neuroinfo/ |
| QuPath | QuPath | RRID: SCR_018257;<br>https://qupath.github.io/ |
| Synapse | TDT | RRID: SCR_006307<br>https://www.synapse.org/ |
| Kilosort | Cortex Lab - UCL | RRID: SCR_016422<br>https://github.com/cortex-lab/KiloSort |
| Phy | Cortex Lab - UCL | N/A https://github.com/cortex-lab/phy |
| IC Capture | The Imaging Source | RRID: SCR_016047<br>https://www.theimagingssource.com/products/software/software-development-kits-sdks/ic-imaging-control/ |
| GraphPad Prism Version 10 | GraphPad | RRID: SCR_002798<br>https://www.graphpad.com/ |
| NWB Conversion Pipeline | CatalystNeuro | https://github.com/catalystneuro/nelson-lab-to-nwb |
| **Other** |  |  |
| 32 Channel Fixed Optrode Array | Innovative Neurophysiology | Custom;<br>http://www.inphysiology.com/optogenetic-applications/ |
| 64 Channel Silicon probe | Cambridge Neurotech | Cat# H6 |
| 400 μm optic fibers | Doric | MFC_400/430-0.66_4mm_MF1.25_FLT |
| 10-Channel Slip Ring Electrical Commutator | Dragonfly | Model SL-10-C;<br>https://campdeninstruments.com/products/10-ch-slip-ring-commuta |
| Master-8 | AMPI | http://www.ampi.co.il/master8cp.html |
| M-Series (M60) Amplifier System for Multiplexing | Triangle BioSystems | http://www.trianglebiosystems.com/m-series-systems.html |
| Single Channel Temperature Controller | Warner Instruments | Cat#TC-324C |
| MINIPULS 3 Peristaltic Pumps | Gilson | Cat#F155008 |
| X-Cite 120LED Boost | Excelitas | Cat#010-00326R |
| ITC-18 16-bit Multi-Channel Data Acquisition Interface | Heka | RRID: SCR_023164 |
| Multiclamp 700B Microelectrode Amplifier | Molecular Devices | RRID: SCR_018455 |
| RHD USB Interface Board | Intan Technologies | Cat# C3100 |
| Assisted Electrical Rotary Joint | Doric | AERJ_24_PZN12 |
| Video camera | The Imaging Source | DMK 33UX273 |

## SUPPLEMENTAL INFORMATION

Document S1. Supplemental Figures S1-S11.

Table S1. Experimental design and statistical analysis of all key experiments

## REFERENCES

1. Goetz, C.G. (2011). The history of Parkinson’s disease: early clinical descriptions and neurological therapies. Cold Spring Harb. Perspect. Med. 1, a008862. 10.1101/cshperspect.a008862.

2. Kalia, L. V, and Lang, A.E. (2015). Parkinson’s disease. Lancet 386, 896–912. 10.1016/S0140-6736(14)61393-3.

3. Carta, M., and Bezard, E. (2011). Contribution of pre-synaptic mechanisms to L-DOPA-induced dyskinesia. Neuroscience 198, 245–251. 10.1016/j.neuroscience.2011.07.070.

4. Connolly, B.S., and Lang, A.E. (2014). Pharmacological treatment of Parkinson disease: a review. JAMA 311, 1670–1683. 10.1001/jama.2014.3654.

5. Cenci, M.A. (2014). Presynaptic Mechanisms of l-DOPA-Induced Dyskinesia: The Findings, the Debate, and the Therapeutic Implications. Front. Neurol. 5, 242. 10.3389/fneur.2014.00242.

6. Jankovic, J. (2005). Motor fluctuations and dyskinesias in Parkinson’s disease: clinical manifestations. Mov. Disord. 20 Suppl 1, S11–6. 10.1002/mds.20458.

7. Cenci, M.A., Riggare, S., Pahwa, R., Eidelberg, D., and Hauser, R.A. (2020). Dyskinesia matters. Mov. Disord. 35, 392–396. 10.1002/mds.27959.

8. Espay, A.J., Morgante, F., Merola, A., Fasano, A., Marsili, L., Fox, S.H., Bezard, E., Picconi, B., Calabresi, P., and Lang, A.E. (2018). Levodopa-induced dyskinesia in Parkinson disease: Current and evolving concepts. Ann. Neurol. 84, 797–811. 10.1002/ana.25364.

9. Manson, A., Stirpe, P., and Schrag, A. (2012). Levodopa-Induced-Dyskinesias Clinical Features, Incidence, Risk Factors, Management and Impact on Quality of Life. J. Parkinsons Dis. 2, 189–198. 10.3233/JPD-2012-120103.

10. Ryan, M.B., Bair-Marshall, C., and Nelson, A.B. (2018). Aberrant Striatal Activity in Parkinsonism and Levodopa-Induced Dyskinesia. Cell Rep. 23, 3438–3446.e5. 10.1016/j.celrep.2018.05.059.

11. Twedell, E.L., Bair-Marshall, C.J., Girasole, A.E., Scaria, L.K., Sridhar, S., and Nelson, A.B. (2024). Striatal lateral inhibition regulates action selection in a mouse model of levodopa-induced dyskinesia. Preprint at bioRxiv, 10.1101/2024.10.11.617939

12. Girasole, A.E., Lum, M.Y., Nathaniel, D., Bair-Marshall, C.J., Guenthner, C.J., Luo, L., Kreitzer, A.C., and Nelson, A.B. (2018). A Subpopulation of Striatal Neurons Mediates Levodopa-Induced Dyskinesia. Neuron 97, 787–795.e6. 10.1016/j.neuron.2018.01.017.

13. Ryan, M.B., Girasole, A.E., Flores, A.J., Twedell, E.L., McGregor, M.M., Brakaj, R., Paletzki, R.F., Hnasko, T.S., Gerfen, C.R., and Nelson, A.B. (2024). Excessive firing of dyskinesia-associated striatal direct pathway neurons is gated by dopamine and excitatory synaptic input. Cell Rep. 43, 114483. 10.1016/j.celrep.2024.114483.

14. Fieblinger, T., Graves, S.M., Sebel, L.E., Alcacer, C., Plotkin, J.L., Gertler, T.S., Chan, C.S., Heiman, M., Greengard, P., Cenci, M.A., et al. (2014). Cell type-specific plasticity of striatal projection neurons in parkinsonism and L-DOPA-induced dyskinesia. Nat. Commun. 5, 5316. 10.1038/ncomms6316.

15. Shen, W., Plotkin, J.L., Francardo, V., Ko, W.K.D., Xie, Z., Li, Q., Fieblinger, T., Wess, J., Neubig, R.R., Lindsley, C.W., et al. (2015). M4 Muscarinic Receptor Signaling Ameliorates Striatal Plasticity Deficits in Models of L-DOPA-Induced Dyskinesia. Neuron 88, 762–773. 10.1016/j.neuron.2015.10.039.

16. Murer, M.G., and Moratalla, R. (2011). Striatal Signaling in L-DOPA-Induced Dyskinesia: Common Mechanisms with Drug Abuse and Long Term Memory Involving D1 Dopamine Receptor Stimulation. Front. Neuroanat. 5, 51. 10.3389/fnana.2011.00051.

17. Paz, R.M., Tubert, C., Stahl, A.M., Amarillo, Y., Rela, L., and Murer, M.G. (2021). Levodopa Causes Striatal Cholinergic Interneuron Burst-Pause Activity in Parkinsonian Mice. Mov. Disord. 10.1002/mds.28516.

18. Sáez, M., Keifman, E., Alberquilla, S., Coll, C., Reig, R., Murer, M.G., and Moratalla, R. (2023). D2 dopamine receptors and the striatopallidal pathway modulate L-DOPA-induced dyskinesia in the mouse. Neurobiol. Dis. 186, 106278. 10.1016/j.nbd.2023.106278.

19. Keifman, E., Ruiz-DeDiego, I., Pafundo, D.E., Paz, R.M., Solís, O., Murer, M.G., and Moratalla, R. (2019). Optostimulation of striatonigral terminals in substantia nigra induces dyskinesia that increases after L-DOPA in a mouse model of Parkinson’s disease. Br. J. Pharmacol. 176, 2146–2161. 10.1111/bph.14663.

20. Singh, A., and Papa, S.M. (2020). Striatal Oscillations in Parkinsonian Non-Human Primates. Neuroscience 449, 116–122. 10.1016/j.neuroscience.2020.09.004.

21. Pagano, G., Yousaf, T., and Politis, M. (2017). PET Molecular Imaging Research of Levodopa-Induced Dyskinesias in Parkinson’s Disease. Curr. Neurol. Neurosci. Rep. 17, 90. 10.1007/s11910-017-0794-2.

22. Zhai, S., Cui, Q., Wokosin, D., Sun, L., Tkatch, T., Crittenden, J.R., Graybiel, A.M., and Surmeier, D.J. (2025). State-dependent modulation of spiny projection neurons controls levodopa-induced dyskinesia in a mouse model of Parkinson’s disease. Sci. Adv. 11. 10.1126/sciadv.adv8224.

23. Shen, W., Zhai, S., Francardo, V., Cui, Q., Sun, L., Xie, Z., Tkatch, T., Corredor, V.E., Lein, E.S., Ting, J.T., et al. (2026). Cell- and state-specific plasticity of striatal glutamatergic synapses is critical to the expression of levodopa-induced dyskinesia. Neuron 114, 2006–2019.e3. 10.1016/j.neuron.2026.04.023.

24. Gerfen, C.R., Engber, T.M., Mahan, L.C., Susel, Z., Chase, T.N., Monsma, F.J.J., and Sibley, D.R. (1990). D1 and D2 dopamine receptor-regulated gene expression of striatonigral and striatopallidal neurons. Science 250, 1429–1432. 10.1126/science.2147780.

25. Cui, G., Jun, S.B., Jin, X., Pham, M.D., Vogel, S.S., Lovinger, D.M., and Costa, R.M. (2013). Concurrent activation of striatal direct and indirect pathways during action initiation. Nature 494, 238–242. 10.1038/nature11846.

26. Jin, X., and Costa, R.M. (2010). Start/stop signals emerge in nigrostriatal circuits during sequence learning. Nature 466, 457–462. 10.1038/nature09263.

27. Tecuapetla, F., Jin, X., Lima, S.Q., and Costa, R.M. (2016). Complementary Contributions of Striatal Projection Pathways to Action Initiation and Execution. Cell 166, 703–715. 10.1016/j.cell.2016.06.032.

28. Parker, J.G., Marshall, J.D., Ahanonu, B., Wu, Y.-W., Kim, T.H., Grewe, B.F., Zhang, Y., Li, J.Z., Ding, J.B., Ehlers, M.D., et al. (2018). Diametric neural ensemble dynamics in parkinsonian and dyskinetic states. Nature 557, 177–182. 10.1038/s41586-018-0090-6.

29. Alcacer, C., Klaus, A., Mendonça, M., Abalde, S.F., Cenci, M.A., and Costa, R.M. (2025). Abnormal hyperactivity of specific striatal ensembles encodes distinct dyskinetic behaviors revealed by high-resolution clustering. Cell Rep. 44, 115988. 10.1016/j.celrep.2025.115988.

30. Alcacer, C., Andreoli, L., Sebastianutto, I., Jakobsson, J., Fieblinger, T., and Cenci, M.A. (2017). Chemogenetic stimulation of striatal projection neurons modulates responses to Parkinson’s disease therapy. J. Clin. Invest. 127, 720–734. 10.1172/JCI90132.

31. Castela, I., Casado-Polanco, R., Rubio, Y.V.-W., da Silva, J.A., Marquez, R., Pro, B., Moratalla, R., Redgrave, P., Costa, R.M., Obeso, J., et al. (2023). Selective activation of striatal indirect pathway suppresses levodopa induced-dyskinesias. Neurobiol. Dis. 176, 105930. 10.1016/j.nbd.2022.105930.

32. Perez, X.A., Zhang, D., Bordia, T., and Quik, M. (2017). Striatal D1 medium spiny neuron activation induces dyskinesias in parkinsonian mice. Movement Disorders 32, 538–548. 10.1002/mds.26955.

33. F. Hernández, L., Castela, I., Ruiz-DeDiego, I., Obeso, J.A., and Moratalla, R. (2017). Striatal activation by optogenetics induces dyskinesias in the 6-hydroxydopamine rat model of Parkinson disease. Movement Disorders 32, 530–537. 10.1002/mds.26947.

34. Lahiri, A.K., and Bevan, M.D. (2020). Dopaminergic Transmission Rapidly and Persistently Enhances Excitability of D1 Receptor-Expressing Striatal Projection Neurons. Neuron 106, 277–290.e6. 10.1016/j.neuron.2020.01.028.

35. Tsuboi, D., Otsuka, T., Shimomura, T., Faruk, M.O., Yamahashi, Y., Amano, M., Funahashi, Y., Kuroda, K., Nishioka, T., Kobayashi, K., et al. (2022). Dopamine drives neuronal excitability via KCNQ channel phosphorylation for reward behavior. Cell Rep. 40, 111309. 10.1016/j.celrep.2022.111309.

36. Shen, W., Flajolet, M., Greengard, P., and Surmeier, D.J. (2008). Dichotomous Dopaminergic Control of Striatal Synaptic Plasticity. Science (1979). 321, 848–851. 10.1126/science.1160575.

37. Costa, R.M. (2007). Plastic Corticostriatal Circuits for Action Learning. Ann. N. Y. Acad. Sci. 1104, 172–191. 10.1196/annals.1390.015.

38. Calabresi, P., Picconi, B., Tozzi, A., and Di Filippo, M. (2007). Dopamine-mediated regulation of corticostriatal synaptic plasticity. Trends Neurosci. 30, 211–219. 10.1016/j.tins.2007.03.001.

39. Wang, Z., Kai, L., Day, M., Ronesi, J., Yin, H.H., Ding, J., Tkatch, T., Lovinger, D.M., and Surmeier, D.J. (2006). Dopaminergic control of corticostriatal long-term synaptic depression in medium spiny neurons is mediated by cholinergic interneurons. Neuron 50, 443–452. 10.1016/j.neuron.2006.04.010.

40. Reynolds, J.N., Hyland, B.I., and Wickens, J.R. (2001). A cellular mechanism of reward-related learning. Nature 413, 67–70. 10.1038/35092560.

41. Druart, M., Yang, Y.C., Tritsch, N.X., and Sippy, T. (2026). The Cellular and Synaptic Actions of Dopamine During Behavior. Preprint at bioRxiv, 10.64898/2026.08.05.742858

42. Escande, M. V, Taravini, I.R.E., Zold, C.L., Belforte, J.E., and Murer, M.G. (2016). Loss of Homeostasis in the Direct Pathway in a Mouse Model of Asymptomatic Parkinson’s Disease. J. Neurosci. 36, 5686–5698. 10.1523/JNEUROSCI.0492-15.2016.

43. de la Fuente-Fernandez, R. (2004). Levodopa-induced changes in synaptic dopamine levels increase with progression of Parkinson’s disease: implications for dyskinesias. Brain 127, 2747–2754. 10.1093/brain/awh290.

44. Sellnow, R.C., Newman, J.H., Chambers, N., West, A.R., Steece-Collier, K., Sandoval, I.M., Benskey, M.J., Bishop, C., and Manfredsson, F.P. (2019). Regulation of dopamine neurotransmission from serotonergic neurons by ectopic expression of the dopamine D2 autoreceptor blocks levodopa-induced dyskinesia. Acta Neuropathol. Commun. 7, 8. 10.1186/s40478-018-0653-7.

45. Fieblinger, T., Zanetti, L., Sebastianutto, I., Breger, L.S., Quintino, L., Lockowandt, M., Lundberg, C., and Cenci, M.A. (2018). Striatonigral neurons divide into two distinct morphological-physiological phenotypes after chronic L-DOPA treatment in parkinsonian rats. Sci. Rep. 8, 10068. 10.1038/s41598-018-28273-5.

46. Gomez, G., Escande, M. V., Suarez, L.M., Rela, L., Belforte, J.E., Moratalla, R., Murer, M.G., Gershanik, O.S., and Taravini, I.R.E. (2019). Changes in Dendritic Spine Density and Inhibitory Perisomatic Connectivity onto Medium Spiny Neurons in l-Dopa-Induced Dyskinesia. Mol. Neurobiol. 56, 6261–6275. 10.1007/s12035-019-1515-4.

47. Suárez, L.M., Solís, O., Caramés, J.M., Taravini, I.R., Solís, J.M., Murer, M.G., and Moratalla, R. (2014). L-DOPA treatment selectively restores spine density in dopamine receptor D2-expressing projection neurons in dyskinetic mice. Biol. Psychiatry 75, 711–722. 10.1016/j.biopsych.2013.05.006.

48. Picconi, B., Centonze, D., Håkansson, K., Bernardi, G., Greengard, P., Fisone, G., Cenci, M.A., and Calabresi, P. (2003). Loss of bidirectional striatal synaptic plasticity in L-DOPA-induced dyskinesia. Nat. Neurosci. 6, 501–506. 10.1038/nn1040.

49. Cenci, M.A., and Lundblad, M. (2007). Ratings of L-DOPA-Induced Dyskinesia in the Unilateral 6-OHDA Lesion Model of Parkinson’s Disease in Rats and Mice. Curr. Protoc. Neurosci. 41. 10.1002/0471142301.ns0925s41.

50. Santini, E., Alcacer, C., Cacciatore, S., Heiman, M., Hervé, D., Greengard, P., Girault, J.-A., Valjent, E., and Fisone, G. (2009). L-DOPA activates ERK signaling and phosphorylates histone H3 in the striatonigral medium spiny neurons of hemiparkinsonian mice. J. Neurochem. 108, 621–633. 10.1111/j.1471-4159.2008.05831.x.

51. Cenci, M.A., Jörntell, H., and Petersson, P. (2018). On the neuronal circuitry mediating L-DOPA-induced dyskinesia. J. Neural Transm. (Vienna) 125, 1157–1169. 10.1007/S00702-018-1886-0.

52. Berton, O., Guigoni, C., Li, Q., Bioulac, B.H., Aubert, I., Gross, C.E., DiLeone, R.J., Nestler, E.J., and Bezard, E. (2009). Striatal Overexpression of ΔJunD Resets L-DOPA-Induced Dyskinesia in a Primate Model of Parkinson Disease. Biol. Psychiatry 66, 554–561. 10.1016/j.biopsych.2009.04.005.

53. Pavón, N., Martín, A.B., Mendialdua, A., and Moratalla, R. (2006). ERK phosphorylation and FosB expression are associated with L-DOPA-induced dyskinesia in hemiparkinsonian mice. Biol. Psychiatry 59, 64–74. 10.1016/j.biopsych.2005.05.044.

54. Santini, E., Valjent, E., Usiello, A., Carta, M., Borgkvist, A., Girault, J.-A., Hervé, D., Greengard, P., and Fisone, G. (2007). Critical involvement of cAMP/DARPP-32 and extracellular signal-regulated protein kinase signaling in L-DOPA-induced dyskinesia. J. Neurosci. 27, 6995–7005. 10.1523/JNEUROSCI.0852-07.2007.

55. Gage, G.J., Stoetzner, C.R., Wiltschko, A.B., and Berke, J.D. (2010). Selective Activation of Striatal Fast-Spiking Interneurons during Choice Execution. Neuron 67, 466–479. 10.1016/j.neuron.2010.06.034.

56. Harris, K.D., Henze, D.A., Csicsvari, J., Hirase, H., and Buzsáki, G. (2000). Accuracy of Tetrode Spike Separation as Determined by Simultaneous Intracellular and Extracellular Measurements. J. Neurophysiol. 84, 401–414. 10.1152/jn.2000.84.1.401.

57. Berke, J.D., Okatan, M., Skurski, J., and Eichenbaum, H.B. (2004). Oscillatory Entrainment of Striatal Neurons in Freely Moving Rats. Neuron 43, 883–896. 10.1016/j.neuron.2004.08.035.

58. Sun, F., Zhou, J., Dai, B., Qian, T., Zeng, J., Li, X., Zhuo, Y., Zhang, Y., Wang, Y., Qian, C., et al. (2020). Next-generation GRAB sensors for monitoring dopaminergic activity in vivo. Nat. Methods 17, 1156–1166. 10.1038/s41592-020-00981-9.

59. Wickens, J.R., Reynolds, J.N., and Hyland, B.I. (2003). Neural mechanisms of reward-related motor learning. Curr. Opin. Neurobiol. 13, 685–690. 10.1016/j.conb.2003.10.013.

60. Surmeier, D.J., Bargas, J., Hemmings, H.C., Nairn, A.C., and Greengard, P. (1995). Modulation of calcium currents by a D1 dopaminergic protein kinase/phosphatase cascade in rat neostriatal neurons. Neuron 14, 385–397. 10.1016/0896-6273(95)90294-5.

61. Shen, W., Hernandez-Lopez, S., Tkatch, T., Held, J.E., and Surmeier, D.J. (2004). Kv1.2-containing K+ channels regulate subthreshold excitability of striatal medium spiny neurons. J. Neurophysiol. 91, 1337–1349. 10.1152/jn.00414.2003.

62. Nisenbaum, E.S., Xu, Z.C., and Wilson, C.J. (1994). Contribution of a slowly inactivating potassium current to the transition to firing of neostriatal spiny projection neurons. J. Neurophysiol. 71, 1174–1189. 10.1152/jn.1994.71.3.1174.

63. Pineda, J.C., Galarraga, E., Bargas, J., Cristancho, M., and Aceves, J. (1992). Charybdotoxin and apamin sensitivity of the calcium-dependent repolarization and the afterhyperpolarization in neostriatal neurons. J. Neurophysiol. 68, 287–294. 10.1152/jn.1992.68.1.287.

64. Pérez-Garci, E., Bargas, J., and Galarraga, E. (2003). The role of Ca2+ channels in the repetitive firing of striatal projection neurons. Neuroreport 14, 1253–1256. 10.1097/00001756-200307010-00013.

65. Vilchis, C., Bargas, J., Ayala, G.X., Galván, E., and Galarraga, E. (1999). Ca2+ channels that activate Ca2+-dependent K+ currents in neostriatal neurons. Neuroscience 95, 745–752. 10.1016/S0306-4522(99)00493-5.

66. Shen, W., Tian, X., Day, M., Ulrich, S., Tkatch, T., Nathanson, N.M., and Surmeier, D.J. (2007). Cholinergic modulation of Kir2 channels selectively elevates dendritic excitability in striatopallidal neurons. Nat. Neurosci. 10, 1458–1466. 10.1038/nn1972.

67. Marcott, P.F., Mamaligas, A.A., and Ford, C.P. (2014). Phasic Dopamine Release Drives Rapid Activation of Striatal D2-Receptors. Neuron 84, 164–176. 10.1016/j.neuron.2014.08.058.

68. Gong, S., Fayette, N., Heinsbroek, J.A., and Ford, C.P. (2021). Cocaine shifts dopamine D2 receptor sensitivity to gate conditioned behaviors. Neuron 109, 3421–3435.e5. 10.1016/j.neuron.2021.08.012.

69. Darmopil, S., Martín, A.B., De Diego, I.R., Ares, S., and Moratalla, R. (2009). Genetic inactivation of dopamine D1 but not D2 receptors inhibits L-DOPA-induced dyskinesia and histone activation. Biol. Psychiatry 66, 603–613. 10.1016/j.biopsych.2009.04.025.

70. Westin, J.E., Andersson, M., Lundblad, M., and Cenci, M.A. (2001). Persistent changes in striatal gene expression induced by long-term l-DOPA treatment in a rat model of Parkinson’s disease. European Journal of Neuroscience 14, 1171–1176. 10.1046/j.0953-816x.2001.01743.x.

71. Park, J., Kang, S., Lee, Y., Choi, J.-W., and Oh, Y.-S. (2024). Continuous long-range measurement of tonic dopamine with advanced FSCV for pharmacodynamic analysis of levodopa-induced dyskinesia in Parkinson’s disease. Front. Bioeng. Biotechnol. 12. 10.3389/fbioe.2024.1335474.

72. Vu, M.-A.T., Brown, E.H., Wen, M.J., Noggle, C.A., Zhang, Z., Monk, K.J., Bouabid, S., Mroz, L., Graham, B.M., Zhuo, Y., et al. (2024). Targeted micro-fiber arrays for measuring and manipulating localized multi-scale neural dynamics over large, deep brain volumes during behavior. Neuron 112, 909–923.e9. 10.1016/j.neuron.2023.12.011.

73. Patriarchi, T., Cho, J.R., Merten, K., Howe, M.W., Marley, A., Xiong, W.-H., Folk, R.W., Broussard, G.J., Liang, R., Jang, M.J., et al. (2018). Ultrafast neuronal imaging of dopamine dynamics with designed genetically encoded sensors. Science (1979). 360. 10.1126/science.aat4422.

74. Planert, H., Berger, T.K., and Silberberg, G. (2013). Membrane Properties of Striatal Direct and Indirect Pathway Neurons in Mouse and Rat Slices and Their Modulation by Dopamine. PLoS One 8, e57054. 10.1371/journal.pone.0057054.

75. Gertler, T.S., Chan, C.S., and Surmeier, D.J. (2008). Dichotomous anatomical properties of adult striatal medium spiny neurons. J. Neurosci. 28, 10814–10824. 10.1523/JNEUROSCI.2660-08.2008.

76. Jin, X., and Costa, R.M. (2015). Shaping action sequences in basal ganglia circuits. Curr. Opin. Neurobiol. 33, 188–196. 10.1016/j.conb.2015.06.011.

77. Legaria, A.A., Matikainen-Ankney, B.A., Yang, B., Ahanonu, B., Licholai, J.A., Parker, J.G., and Kravitz, A. V. (2022). Fiber photometry in striatum reflects primarily nonsomatic changes in calcium. Nat. Neurosci. 25, 1124–1128. 10.1038/s41593-022-01152-z.

78. Tepper, J.M., Koós, T., and Wilson, C.J. (2004). GABAergic microcircuits in the neostriatum. Trends Neurosci. 27, 662–669. 10.1016/j.tins.2004.08.007.

79. Tepper, J.M., Koós, T., Ibanez-Sandoval, O., Tecuapetla, F., Faust, T.W., and Assous, M. (2018). Heterogeneity and Diversity of Striatal GABAergic Interneurons: Update 2018. Front. Neuroanat. 12, 91. 10.3389/fnana.2018.00091.

80. Taverna, S., Ilijic, E., and Surmeier, D.J. (2008). Recurrent Collateral Connections of Striatal Medium Spiny Neurons Are Disrupted in Models of Parkinson’s Disease. The Journal of Neuroscience 28, 5504–5512. 10.1523/JNEUROSCI.5493-07.2008.

81. Goggi, J.L., Sardini, A., Egerton, A., Strange, P.G., and Grasby, P.M. (2007). Agonist-dependent internalization of D2 receptors: Imaging quantification by confocal microscopy. Synapse 61, 231–241. 10.1002/syn.20360.

82. Sage, J.I., and Mark, M.H. (1994). Basic mechanisms of motor fluctuations. Neurology 44, S10–4.

