## Supplementary Figures and Text for "Repetitive Levodopa Treatment Drives Cell Type-Specific Striatal Adaptations Associated with Progressive Dyskinesia in Parkinsonian Mice"

**SUPPLEMENTAL INFORMATION**

**Figure S1**

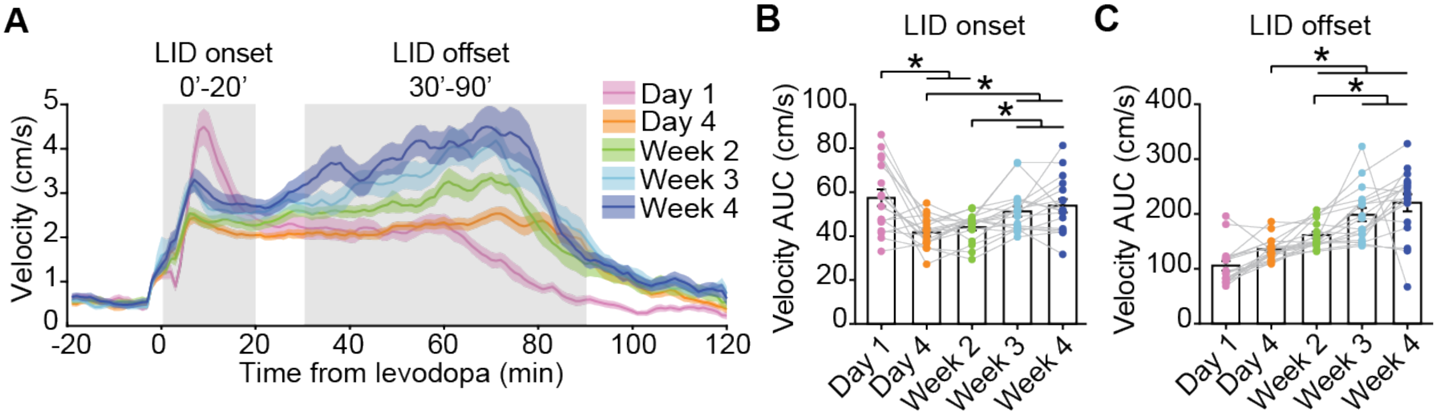


**Figure S1: Changes in locomotor velocity after repetitive levodopa, related to Figure 1. (A)** Average locomotor velocity in response to IP injection of 5mg/kg levodopa for different treatment timepoints. **(B-C)** Area under the curve (AUC) of velocity traces during LID onset (B), and offset (C) phases. [RM one-way ANOVA, B: $p=0.0004$, post-hoc $*p<0.05$; C: $p<0.0001$, post-hoc $*p<0.05$]. Data shown as mean ± SEM. N=17. N=mice.

**Figure S2**

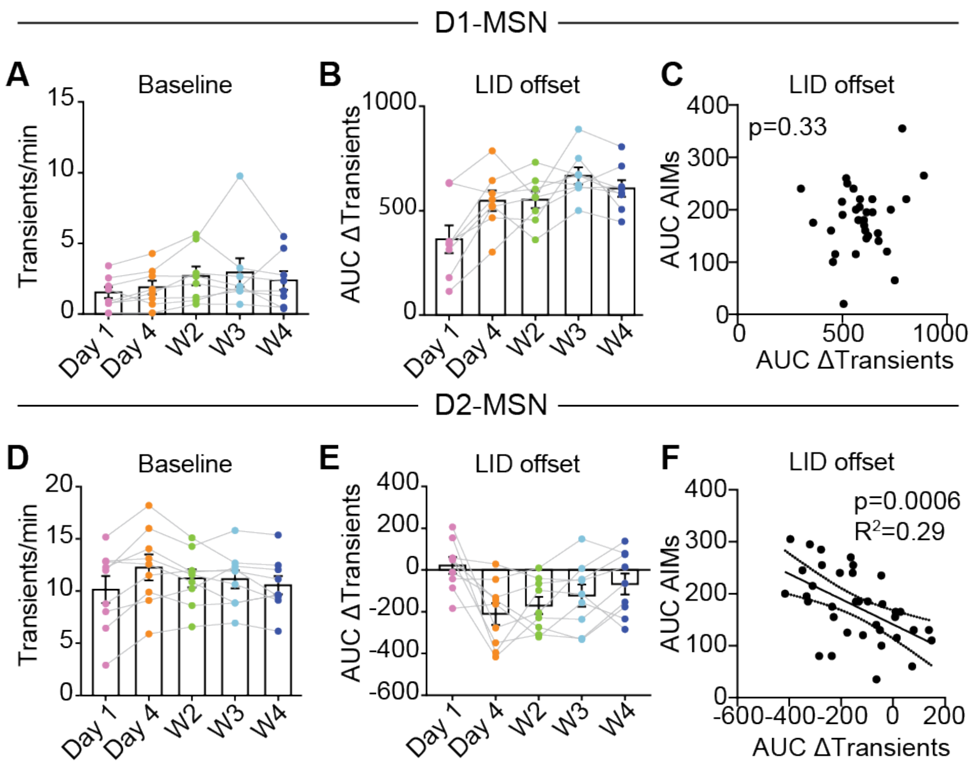


**Figure S2: More rapid LID offset relates to the recovery of D2-MSN activity, related to Figure 2. (A)** Average baseline rates of D1-MSN GCaMP transients 30 min before levodopa injection. [RM one-way ANOVA, p=0.2270]. **(B)** Area under the curve of D1-MSN ΔTransients during LID offset. [RM one-way ANOVA, ns]. **(C)** Correlation of D1-MSN ΔTransients with AIMs during LID offset. **(D)** Average baseline rates of D2-MSN GCaMP transients 30 min before levodopa injection. [RM one-way ANOVA, p=0.1573]. **(E)** Area under the curve of D2-MSN ΔTransients during LID offset. [RM one-way ANOVA, p=0.0436]. **(F)** Correlation of D2-MSN ΔTransients with AIMs during LID offset. N=8 D1-Cre and N=9 A2a-Cre mice. Each dot is one mouse. Data shown as mean ± SEM (A, B, D, E). N = mice.

### Figure S3

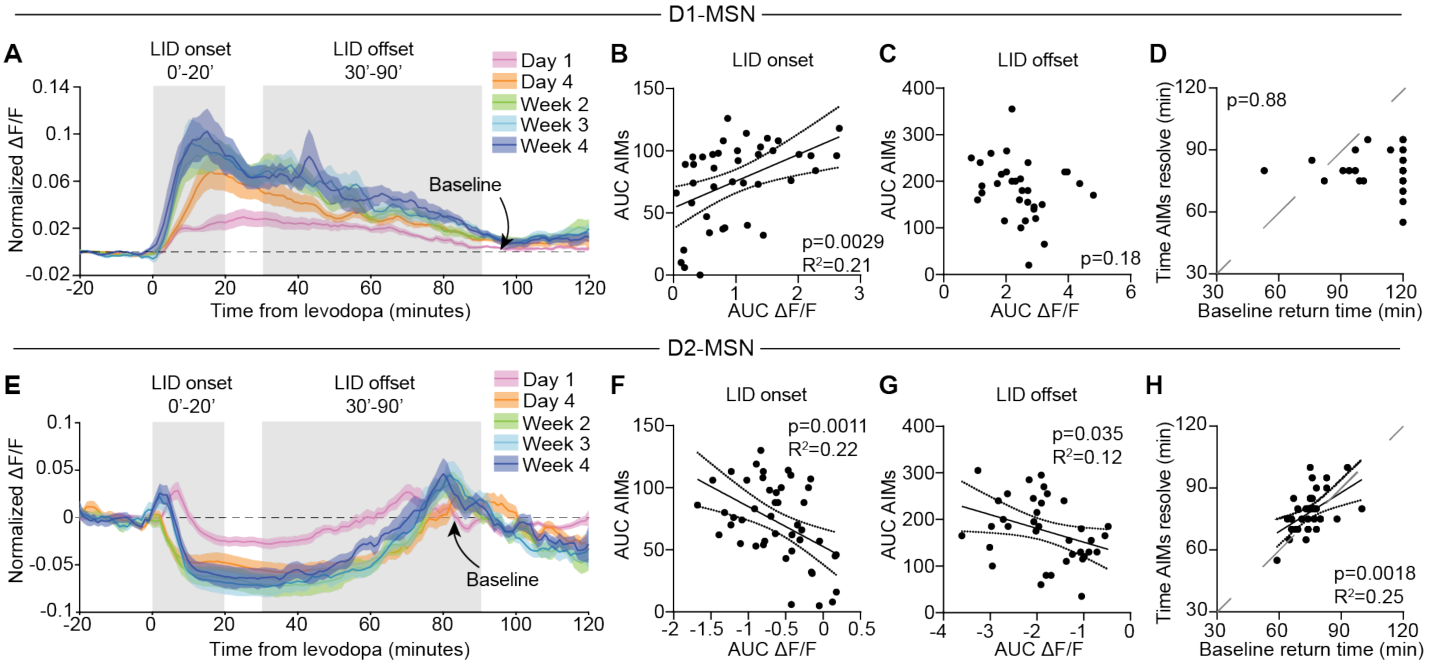


**Figure S3: Changes in striatal activity parallel alterations in dyskinesia during repetitive levodopa treatment, related to Figure 2.** GCaMP fiber photometry in DLS D1-MSNs (A-D) or D2-MSNs (E-H) of 6-OHDA mice treated with levodopa. The readout for MSN activity readout is ΔF/F, rather than transient frequency (Figure 2). **(A)** D1-MSN ΔF/F response to levodopa with baseline subtraction across treatment timepoints. LID onset and offset phases are shown in gray. **(B-C)** Correlation between AIMs and D1-MSN ΔF/F during LID onset (B) and offset (C) phases. **(D)** Correlation between time of AIMs resolution and the time D1-MSN ΔF/F returns to baseline. **(E)** D2-MSN ΔF/F response to levodopa with baseline subtraction across treatment timepoints. **(F-G)** Correlation between AIMs and D2-MSN ΔF/F during LID onset (F) and offset (G) phases. **(H)** Correlation between time of AIMs resolution and the time D2-MSN ΔF/F returns to baseline. N=9 A2a-Cre and N=8 D1-Cre mice. Data shown as mean ± SEM (A, E). N=mice.

### Figure S4

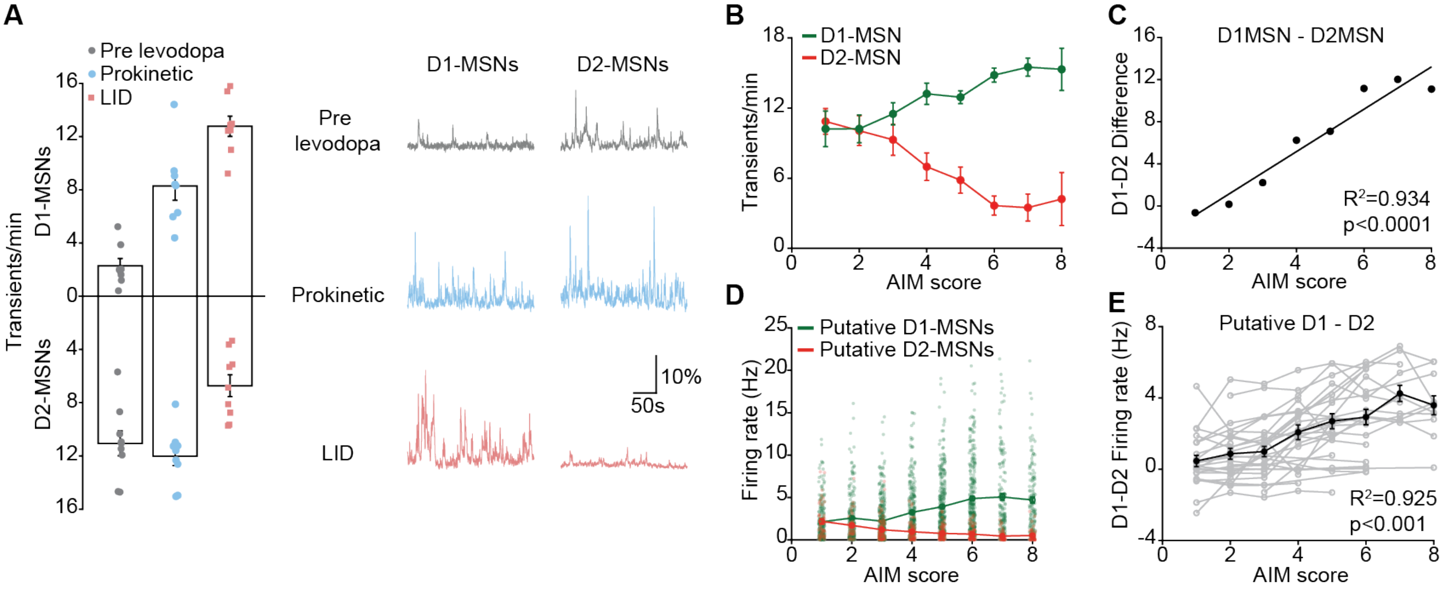


**Figure S4: Relative D1/D2-MSN activity relates to prokinetic effects and dyskinesia severity, related to Figures 2 & 4. (A)** Average GCaMP6s transient frequency in D1- and D2-MSNs before levodopa administration (pre-levodopa), during the prokinetic phase (velocity > 1.5 cm/s; AIMs = 0), and during LID (AIMs > 1). Each dot represents the average from all sessions from each mouse. Right: representative GCaMP6s traces from D1- and D2-MSNs. **(B)** Average GCaMP6s transient frequency in D1- and D2-MSNs plotted as a function of LID severity [N=8 D1-Cre and N=9 A2a-Cre mice]. **(C)** Difference between the D1- and D2-MSN traces shown in (B), with linear correlation analysis. **(D)** Average firing rate of putative D1- and D2-MSNs plotted as a function of LID severity (n=372 putative D1-MSNs and n=123 putative D2-MSNs). Each dot represents one single unit. **(E)** Difference between the average firing rates of putative D1- and D2-MSNs as a function of LID severity within each animal (gray lines indicate individual mice; black line indicates the group average). Data shown as mean ± SEM (A, B, D, E). n = cells; N = mice.

**Figure S5**

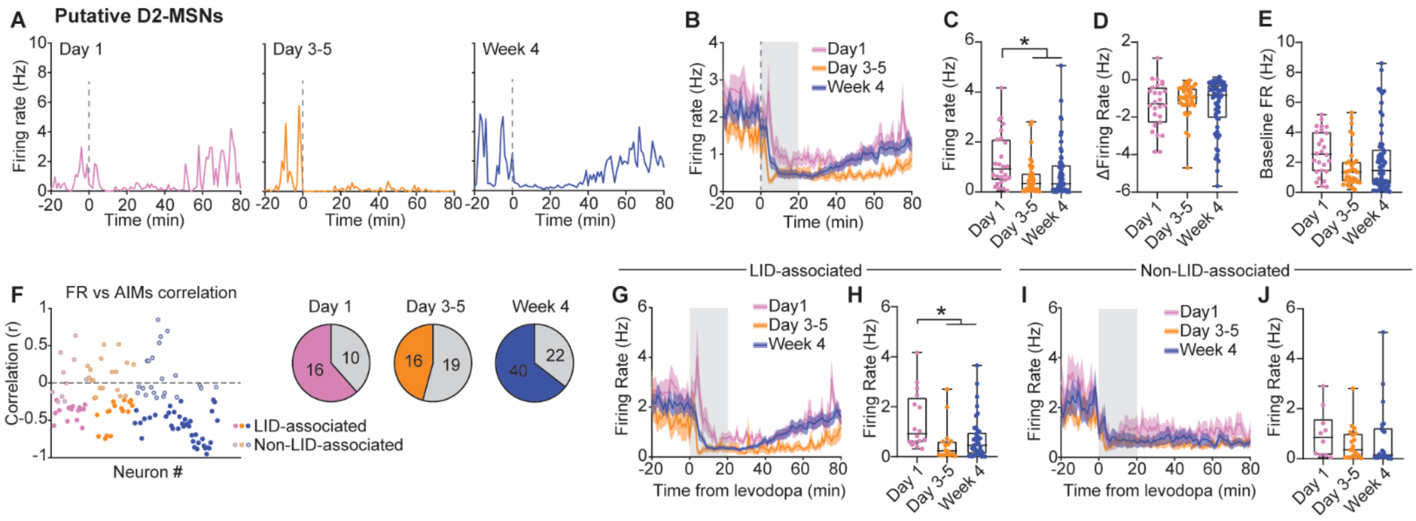


**Figure S5: Repetitive levodopa drives changes in D2-MSN firing rates, related to Figure 4. (A-B)** Representative single units (A) and pooled single unit firing rates (B) of putative D2-MSN on days 1, 3-5 and week 4. **(C-D)** Average firing rate (C) and change in firing rate relative to baseline (D) during LID onset period (gray bar in B) for D2-MSN. [C: Kruskal-Wallis, p=0.0044, post-hoc: *p<0.05; D: Kruskal-Wallis, ns]. **(E)** Average firing rate during baseline (average of 20 minutes pre levodopa) for putative D2-MSNs [Kruskal-Wallis, p=0.0390, post-hoc comparisons: ns]. Day 1: n=26, N=11; Day 4: n=35, N=11; Week 4: n=62, N=15. **(F)** *Left:* Distribution of correlation strengths between the firing rates of putative D2-MSNs and AIM scores across treatment days. Filled dots indicate LID-associated D2-MSNs. *Right:* Proportion of putative D2-MSNs classified as LID-associated (colored) versus non-LID-associated (gray) across treatment days. **(G)** Firing rates of LID-associated D2-MSNs in response to levodopa injections across treatment days. **(H)** Comparison of their average firing rates during the LID onset phase. [Kruskal-Wallis: p=0.0044, post-hoc: *p<0.05]. Day 1: n=16, N=11; Day 4: n=16, N=11; Week 4: n=40, N=15. **(I)** Firing rates of non-LID-associated putative D2-MSNs in response to levodopa injections across treatment days. **(J)** Comparison of their average firing rates during the LID onset phase [Kruskal-Wallis, ns]. Day 1: n=10, N=11; Day 4: n=19, N=11; Week 4: n=22, N=15. n=units; N=animals. Data shown as mean ± SEM (B, G, I) and as median and range (C, D, E, H, J).

**Figure S6**

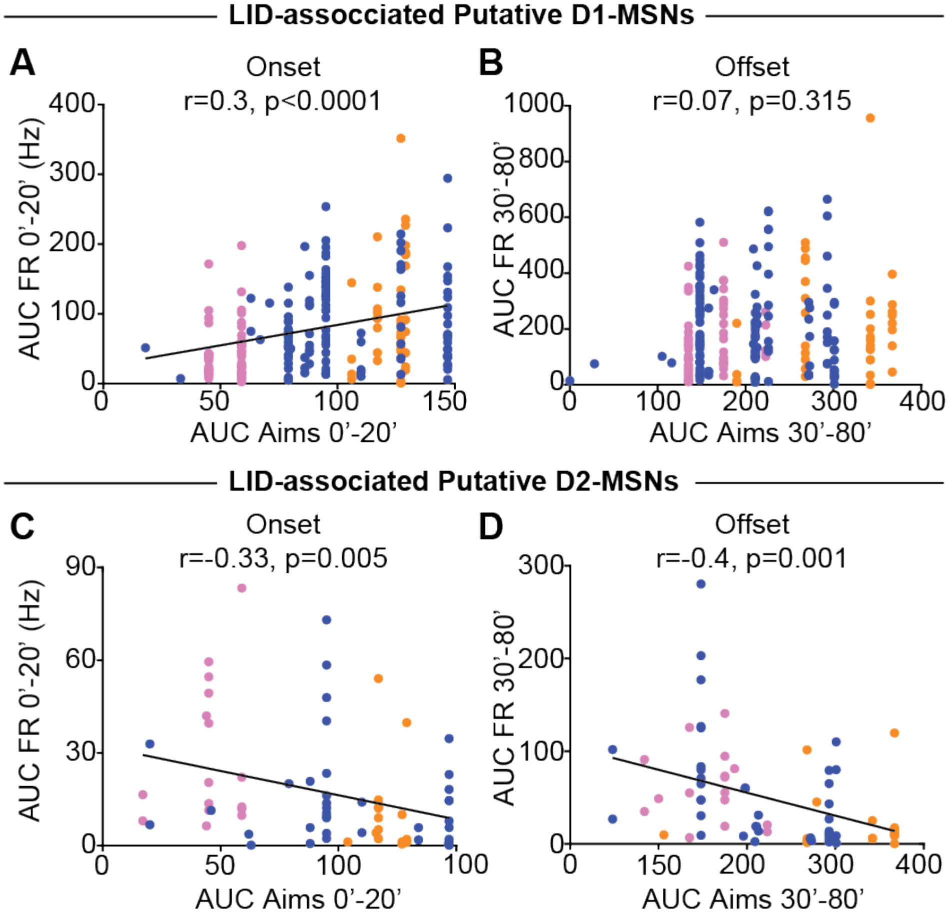


**Figure S6: LID-associated D1- and D2-MSN activity correlates with AIMs during LID onset, but only D2-MSN activity correlates during LID offset, related to Figure 4**. **(A-B)** Correlations of firing rates and AIMs during LID onset (A) and offset (B) periods for LID-associated D1-MSNs. Day 1: n=50, N=11; Day 4: n=38, N=11; Week 4: n=122, N=15. **(C-D)** Correlations of firing rates and AIMs during LID onset (A) and offset (B) periods for LID-associated D2-MSNs. Day 1: n=16, N=11; Day 4: n=16, N=11; Week 4: n=40, N=15. The r and p values of the correlations are indicated on each panel. Color code: pink: units recorded on day 1; orange: days 3-5; and blue: week 4. Each dot represents one unit.

**Figure S7**

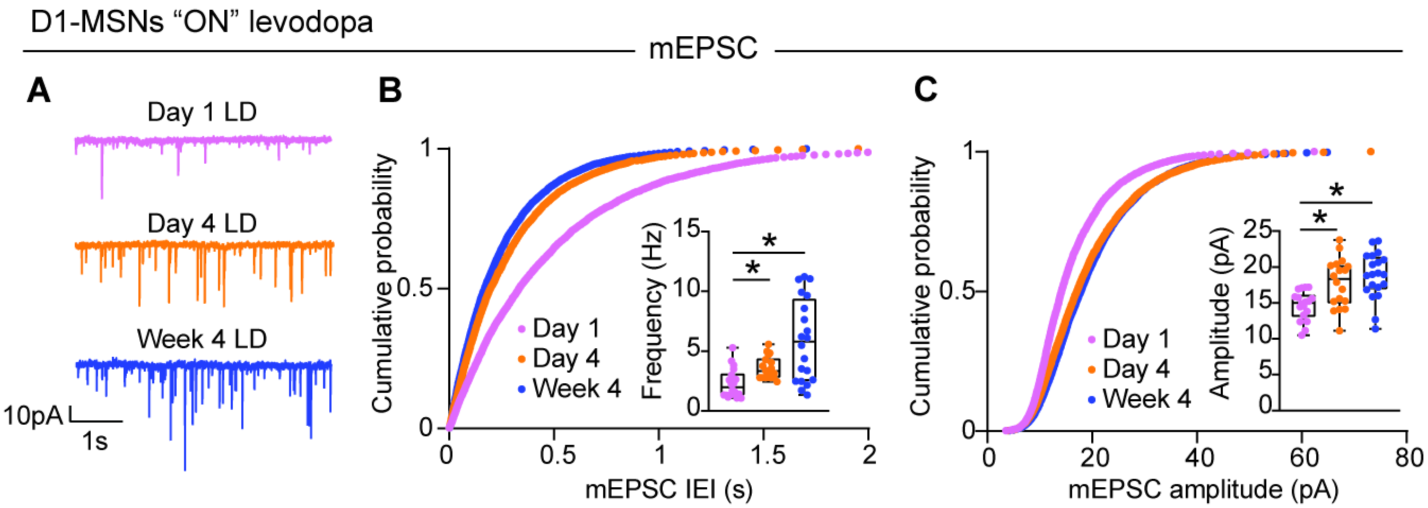


**Figure S7: Repetitive levodopa treatments drive increased excitatory transmission onto D1-MSNs in the "ON" state, related to Figure 6.** 6-OHDA-treated Drd1-tdTomato mice were sacrificed at three different time points for *ex vivo* (slice) electrophysiology, 30 minutes after levodopa injection. Slice recordings were included for up to 4 hours after levodopa administration. **(A)** Representative voltage clamp recordings of miniature excitatory postsynaptic currrents (mEPSC) in D1-MSNs across treatment timepoints. **(B)** Cumulative probability distribution of mEPSC Inter-Event-Intervals (IEI) in D1-MSNs. Inset: average mEPSC frequency. [Kruskal-Wallis, p=0.0003, post-hoc comparisons *p<0.05]. **(C)** Cumulative probability distribution of mEPSC amplitudes in D1-MSNs. Inset: average mEPSC amplitude. [Kruskal-Wallis, p=0.0003, post-hoc comparisons *p<0.05]. Day 1: n=18, N=3; Day 4: n=18, N=3; Week 4: n=20, N=3. n=cells; N=mice. Data shown as mean for probability plots, and median and range for insets.

**Figure S8**

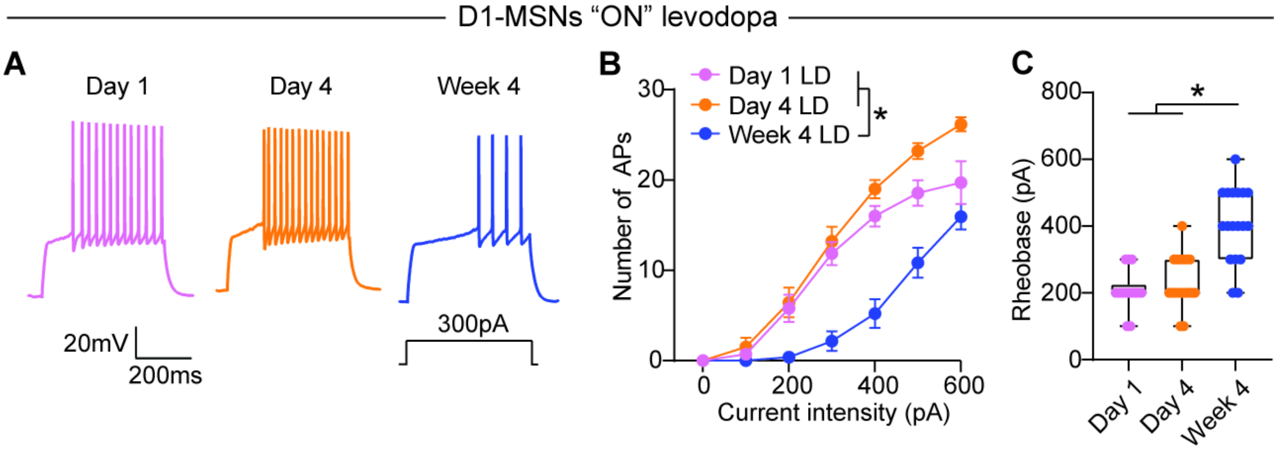


**Figure S8: D1-MSN intrinsic excitability during the "ON" state is reduced after repetitive levodopa treatment, related to Figure 7.** 6-OHDA-treated animals were sacrificed 30 minutes after levodopa injection at three different time points for *ex vivo* (slice) electrophysiology, either after the first dose of levodopa ("Day 1 LD"), after the fourth levodopa administration (Day 4) or during week 4 (Week 4 LD). Slice recordings were included for up to 4 hours after levodopa administration. **(A)** Voltage responses to current injection in D1-MSN after different levodopa treatment duration. **(B)** Number of action potentials (APs) in response to depolarizing current steps. [Two-way ANOVA, interaction $p<0.0001$, post-hoc for 200-500pA $*p<0.05$]. **(C)** D1-MSN rheobase across treatment duration. [Kruskal-Wallis, $p<0.0001$, post-hoc $*p<0.001$]. Day 1 LD: n=14, N=3; Day 4 LD: n=18, N=3; Week 4 LD: n=18, N=3. n=cells; N=mice. Data shown as mean ± SEM (B) and as median and range (C). Each dot represents one cell.

**Figure S9**

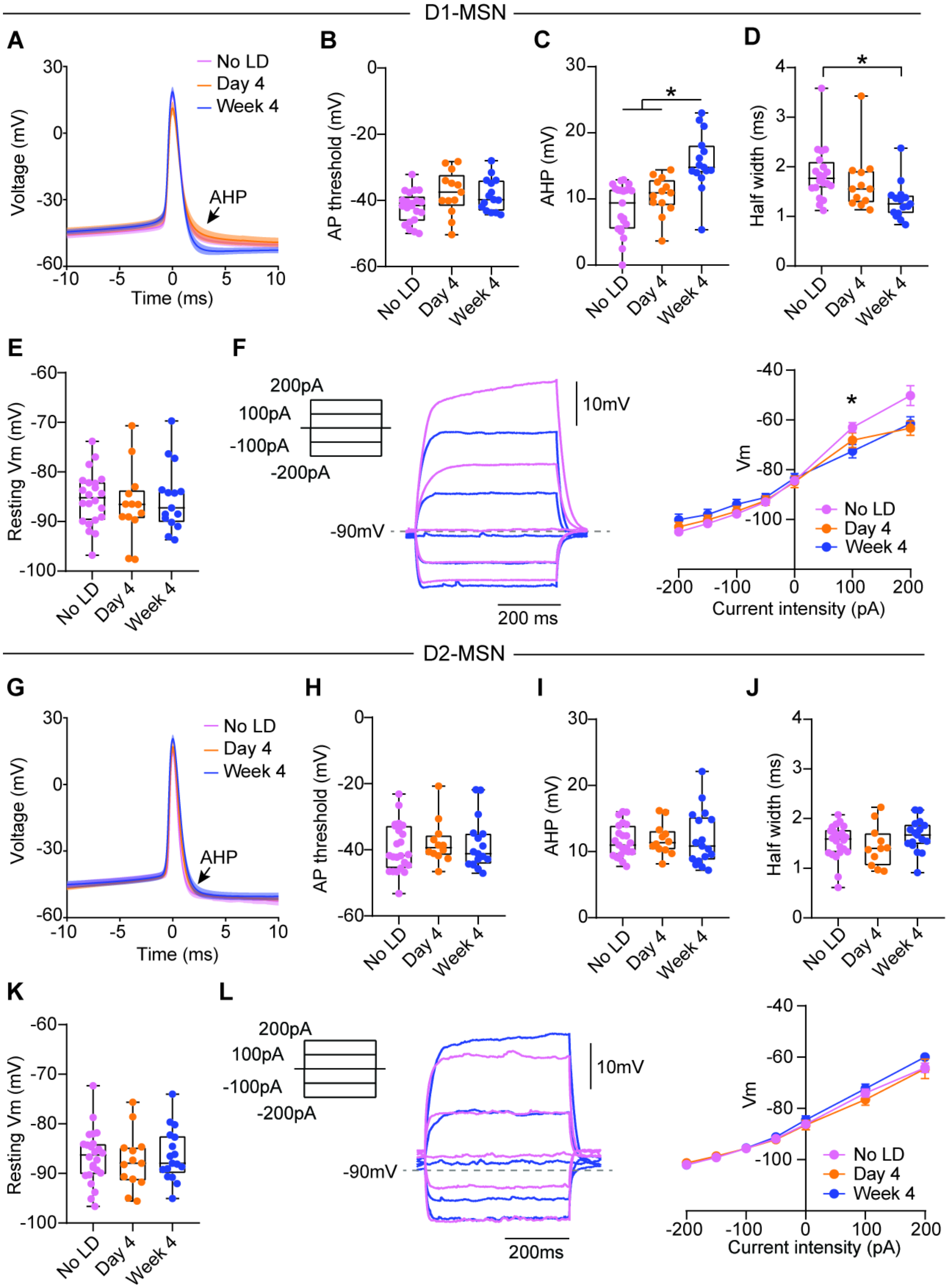


**Figure S9: Changes in intrinsic properties of D1- and D2-MSN after repetitive levodopa measured in the "OFF" state, related to Figure 7. (A)** Average action potential shape for D1-MSNs. **(B-E)** Action potential threshold (B), afterhyperpolarization (AHP, C), half width (D) and resting membrane potential (E) for D1-MSNs. [Kruskal-Wallis, ns (B, E), $p<0.0001$ (C), $p=0.0021$ (D), post-hoc:

*p<0.05]. **(F)** Voltage deflection to current injections in D1-MSNs. [Two-way ANOVA, interaction: $p<0.0001$, post-hoc between No LD and Week 4 at 100pA: *p=0.0211]. No LD: n=20-22, N=9; Day 4: n=12-13, N=6; Week 4: n=15, N=6. **(G)** Average action potential shape for D2-MSNs. **(H-K)** Action potential threshold (H), afterhyperpolarization (AHP, I), half width (J) and resting membrane potential (K) for D2-MSNs. [Kruskal-Wallis, ns]. **(L)** Voltage deflection to current injections in D2-MSNs. [Two-way ANOVA, ns]. No LD: n=22-24, N=11; Day 4: n=11-13, N=6; Week 4: n=17, N=5. Data shown as mean $\pm$ SEM (A, F, G, L) and as median and range (B-E, H-K). n=cells; N=mice. Each dot represents one cell.

**Figure S10**

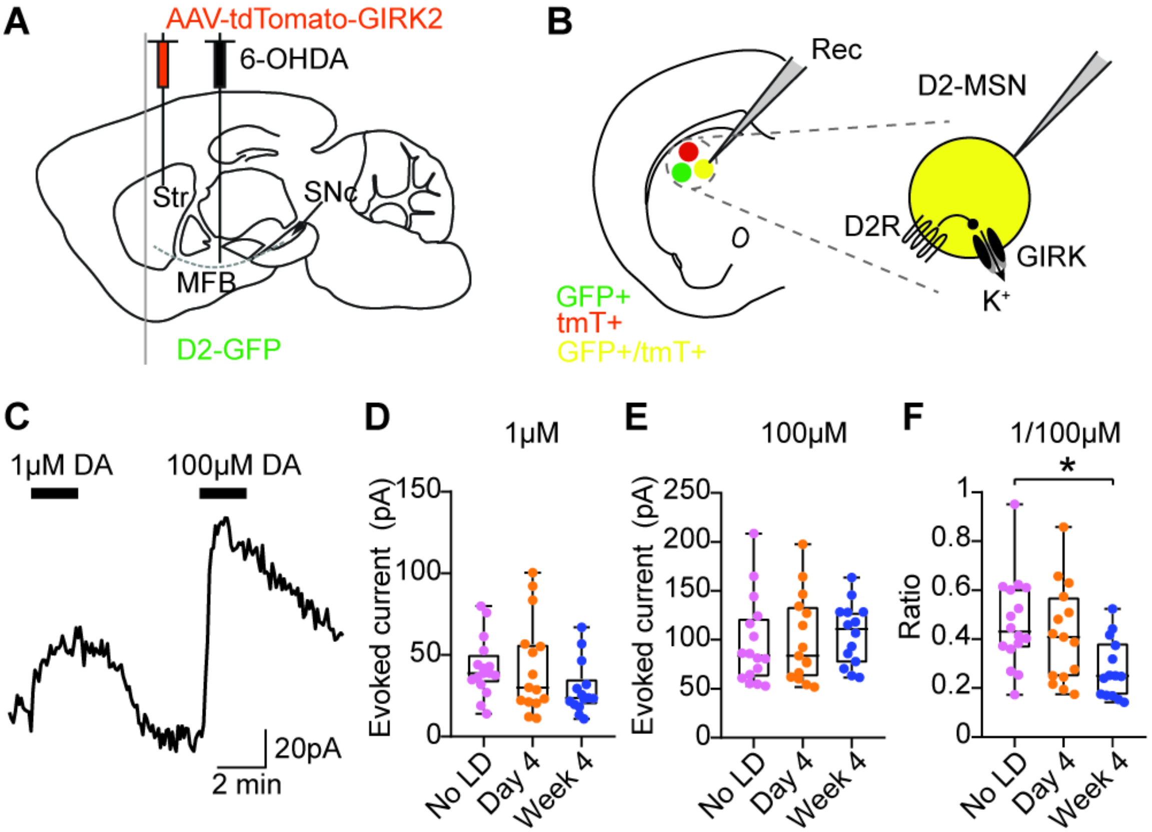


**Figure S10: D2R sensitivity is reduced after 4 weeks of levodopa, related to Figure 8.** D2-GFP mice were injected with an AAV encoding tdTomato-GIRK2 in the dorsolateral striatum, and injected with 6-OHDA in the MFB. **(A)** Experimental approach. **(B)** Schematic showing whole-cell recording of D2-MSNs expressing GIRK (GFP+/tdTomato+ cells). **(C)** Representative current trace showing responses to bath application of 1μM and 100μM dopamine (DA). **(D-F)** GIRK-mediated currents evoked by 1μM (D) or 100μM (E) dopamine and their ratio (F) in D2-MSN after different levodopa treatment timepoints. [F: Kruskal-Wallis, $p=0.01$, post-hoc: $^*p<0.05$]. No LD: n=16, N=7; day 4: n=15, N=7; week 4: n=14, N=8. n=cells; N=mice. Data shown as median and range. Each dot represents one cell.

**Figure S11**

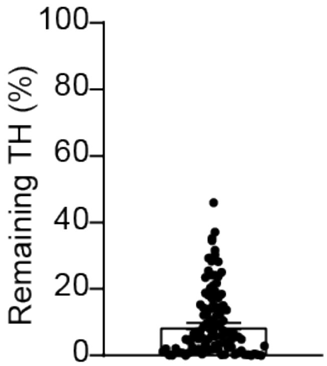


**Figure S11: Average remaining TH immunostaining in 6-OHDA-lesioned striatum relative to the contralateral dopamine-intact side, related to STAR Methods.** Each dot is one mouse. N = 115 mice.
